# HPV integration in head and neck cancer: downstream splicing events and expression ratios linked with poor outcomes

**DOI:** 10.1101/2025.01.17.633627

**Authors:** Shiting Li, Shaomiao Xia, Maria Lawas, Aishani Kulshreshtha, Bailey F. Garb, AA Chamila Perera, Chen Li, Tingting Qin, Joshua D. Welch, Nisha J. D’Silva, Laura S. Rozek, Maureen A. Sartor

## Abstract

HPV integration (HPVint) is associated with carcinogenesis and tumor progression in HPV-associated cancers, including head and neck squamous cell carcinomas (HNSCC). While its impact on human DNA has been well characterized, its relationship with clinical outcomes remains unconfirmed. Here we investigate the consequences of HPVint both with respect to human and HPV characteristics by analyzing 261 HPV-associated HNSCC bulk and single-cell RNA-seq samples from five cohorts, and DNA HPVint events from 102 HPV+ participants in two of the cohorts. By leveraging this large meta-cohort, we first reveal an oncogenic network based on the recurrent HPV integration locations in HNSCC. We then classify HPVint-positive (HPVint(+)) participants by HPV RNA features, specifically based on spliced HPV-human fusion transcripts and ratios of HPV gene transcripts, showing that subsets of participants have worse clinical outcomes. Our analyses, focused mainly on RNA instead of DNA, expand our understanding of the carcinogenic mechanisms of HPVint, partially addressing the conflicting findings of whether HPVint is associated with aggressive phenotypes and worse clinical consequences, and provide potential biomarkers to advance precision oncology in HPV-associated HNSCC.

## Introduction

Human papillomavirus (HPV)-associated head and neck squamous cell carcinoma (HPV(+) HNSCC) has surpassed cervical cancer (CC) as the most common HPV-associated cancer in the U.S.^1^ and global rates of HPV(+) HNSCC are rising.^2,3^ The overall better prognosis of HPV(+) HNSCC patients (∼80% 5-year survival) compared to HPV(-) HNSCC has prompted the exploration of de-escalated treatment regimens for a subset of HPV(+) HNSCC patients^4^, aiming to reduce treatment-related toxicity without compromising efficacy. However, the challenge lies in establishing a clear, reliable criteria for identifying which patients can safely benefit from de-escalated treatment, and conversely which may benefit from more aggressive therapy. HPV integration (HPVint) has been reported to be associated with oncogenic development and progression in both CC and HNSCC ^5,6^ and multiple lines of evidence suggest it as a potential therapeutic target due to its promotion of genomic instability and immune evasion^7^.

However, the downstream effects of HPVint in HNSCC remain incompletely understood with inconsistent findings regarding clinical outcomes ^2,8–11^. Among HPV(+) HNSCC, the estimated proportion of HPVint(+) cases (∼60-70%) ^12,13^ substantially exceeds the percentage exhibiting poor clinical outcomes (∼25-30%) ^14,15^. Consequently, a critical knowledge gap persists regarding the extent to which HPVint affects clinical outcomes in HNSCC and whether only a subset of HPVint(+) HNSCC patients experience worse prognoses.

HPV comprises a double-stranded, approximately 8 kb circular DNA with tightly regulated expression^16^. High-risk HPV initiates infection in the basal layer of the mucosal epithelium^16–18^. Early in its lifecycle, HPV expresses the genes E1 and E2 related to viral replication, along with the viral oncoproteins E6 and E7; as infected keratinocytes differentiate, HPV activates the expression of E4 and E5, followed by the late-expressing capsid and assembly genes, L1 and L2 ^17^. Normally, HPV regulates its gene expression through autoregulation ^18,19^ and splicing ^19–21^. For instance, in HPV16, splice sites located in genes E1, E2 and E4 (relative position: E1:881, E2:2708, and E4: 3357) regulate E2 and E4 expression ^20,22^. HPVint often results in a dramatically perturbation of relative HPV gene expression levels, potentially contributing to oncogenic transformation. For instance, HPVint frequently disrupts E2 ^23–25^, which regulates the transcription of other E genes, leading to uncontrolled expression of E6 and E7 ^26–29^ , further inactivating key tumor suppressor proteins including p53 (*TP53*) and RB transcriptional corepressor 1 (*RB1*), respectively ^30^.

In addition to dysregulation of HPV genes, HPVint in HNSCC and CC are overrepresented in human-specific genomic unstable regions (recurrent regions) ^31,32^, including keratinocyte-specific super-enhancers^33^, highly conserved CTCF binding regions ^6^, cancer-related genes (e.g. MYC/PVT1, MACROD2^12^, DIAPH2, TP63, and NAP1)^34^ and immune-related genes (e.g. PDL1/PDL2/PLGRKT, TNFRSF13B, JAK1) ^26^. HPVint can upregulate oncogenes revealing potential therapeutic targets for HPV-related cancers ^33^. In addition, both innate and adaptive immune responses may be compromised by HPVint in HNSCC ^2^ with HPVint associated with lower levels of T cells, B cells, and NK cells compared to non-integrated tumors^8^.

A comprehensive understanding of the consequences of HPV integration requires both DNA and RNA-based sequencing techniques, with each offering unique insights. DNA-based methods such as whole genome sequencing (WGS) have the potential to detect all insertions and reveal the precise integration sites and structural alterations within the human genome ^35–37^. Complementing this, RNA-based methods such as RNA-seq examine the consequences of HPVint at the transcriptional level. By measuring the expression levels of viral and host transcripts and detecting viral-host fusion transcripts, RNA-seq can reveal how HPV integration affects gene expression ^38^. The dual approach is crucial to understand how genomic integration drives oncogenic processes. As an alternative to direct evidence from sequencing, the E2/E6 or E2/E7 ratios have served as biomarkers for HPVint, with low ratios indicating HPVint and higher ratios being inconclusive although typically interpreted as episomal ^39,40^.

As mentioned, conflicting results have been observed for the association between HPV integration and survival in both CC ^25,38,41,42^ and HNSCC ^2,8–11^. We noticed that studies detecting HPVint status at the RNA level ^8^ were more likely to reveal worse survival than those using DNA detection in HNSCC ^9–11^. Loss of E2 expression has been reported as a predictive biomarker for OPSCC de-escalation trials due to its correlation with worse disease-free survival ^43^. These studies indicated occurrence of HPVint as a potential biomarker of worse clinical outcomes, requiring further stratification and clarification ^7^. Therefore, in this study, we conducted a comprehensive analysis of HPVint in 261 HPV+ HNSCC RNA-seq samples from multiple cohorts, leading to three key conclusions. First, we reveal the most comprehensive list of recurrent HPV integration locations in the human genome in HNSCC which serves as a resource for treatment target discovery efforts. Second, we define two downstream consequences of HPVint based on spliced HPV-human fusion transcripts and ratios of HPV gene expression levels and link these consequences to worse clinical outcomes. Results reveal that keratinization, cell-to-cell junction, and immune infiltration are pathways associated with HPV integration and/or downstream consequences. Third, our study shows that HPVint at the RNA level is more likely to cause alterations in human and HPV gene expression. We propose that HPVint events with detectable viral-host fusion transcripts are more relevant to carcinogenesis than DNA-only HPVint, and encourage future HPVint research to consider RNA.

## Methods

### Overview of bulk and single-cell RNA-seq data

In this study, we used bulk RNA-seq data from 239 HPV(+) HNSCC samples consisting of 236 unique participants from three previous cohorts (UM_FF (n=18; GSE74956), HVC (n=83; EGAD00001004366), and TCGA HNSC-project (n=73)) and a newly introduced cohort UM_FFPE (n=62; EGAxxxxx), plus 3 cell lines (UM-SCC-47, UM-SCC-090, and UM-SCC-154). In addition, we used single-cell RNA-seq (scRNA-seq) data from 22 HPV(+) HNSCC tumors from three studies (GSE181919 ^44^, GSE182227 ^45^, and GSE234933 ^46^). Tumor samples were sequenced from fresh frozen (FF) tissue except UM_FFPE which was from formalin-fixed paraffin-embedded (FFPE) blocks.

Nine samples in UM_FF were from the same 9 patients as ten samples in UM_FFPE. We chose the results from the UM_FF over UM_FFPE for patient-level downstream analysis, as RNA-seq from FF are generally considered more reliable. Overall, this study included 261 HPV(+) HNSCC RNA-seq samples from 249 participants and 3 cell lines.

### Bulk and single-cell RNA-seq preprocessing, HPV integration detection and annotation

Bulk RNA-seq data preprocessing steps including quality control, adapters trimming, reads alignment, and human gene counts calculation were performed using well-established bioinformatics tools and pipelines (See Supplementary Methods). For scRNA-seq data, fastq files were retrieved from SRA by SRA-prefetch and parallel-fastq-dump, except for GSE234933 which was obtained directly from the authors. Cell-level annotations from the original studies were first examined, and HPV(+) samples with at least 100 epithelial cells were selected for HPV integration calling. HPV integration events for bulk RNA-seq were called using two software tools, Survirus ^47^ and CTAT-VIF ^48^, and a custom pipeline to harmonize final results. For scRNA-seq, we customized a new pipeline based on nf-core/viralintegration ^49^ (See Supplementary Methods). For each integration event identified, we filtered out low quality events and classified them as either unspliced or spliced events (See Supplementary Methods).

We used multiple approaches to annotate HPVint events with human genes potentially influenced by the HPV insertion. By examining expression changes in the annotated genes, we selected integrated, expression outlier genes for further analysis (See Supplementary Methods).

### Defining HPV integration status for each participant

To define patient-level HPVint status, we categorized samples/participants into three groups based on the confidence level of their HPVint events (**Supplementary Figure S1A-B,** Supplementary Methods). Primarily, samples/participants with at least one confident HPV integration event (spliced or unspliced) were classified as HPV integration positive (HPVint(+):149). Those with low-quality HPVint events were labeled as HPVint not determined (HPVint-ND), and the remaining were classified as HPVint negative (HPVint(-)). For participants with both FFPE and FF samples, we used their FF results as gold standard.

We reassigned three UM_FFPE samples initially classified as HPVint(-) or HPVint-ND to HPVint(+) because they had at least 2 unfiltered split reads detected by both programs, and reassigned samples with at least 2 split reads detected by one software to HPVint-ND (**Supplementary Figure S1B,** See Supplementary methods). Because two of these UM_FFPE samples had a matching FF sample already assigned as HPVint(+), at the participant level, only one participant was reassigned to HPVint(+). We also reassigned one sample/participant to HPVint(+) with a HPV L1 ratio score Z-score smaller than -3, since this indicated loss of L1 in this sample (**Supplementary Figure S1C,** See Supplementary methods).

Finally, we stratified the HPVint(+) participants based on the presence of spliced integration events. For participants with E1-related spliced integration events (i.e. splice sites at HPV16 & 35: 880-884, HPV33: 894, HPV18:930) or E6*-related spliced integration events (i.e. splice sites at HPV16: 226-230, 406-408. HPV33: 233), we classified them as having E1* integration or E6* integration, respectively.

### Demographic information collection and statistical testing

We collected demographic information from published papers ^44–46^, TCGA GDC portal, and the UM Head and Neck SPORE database, focusing primarily on gender, self-reported race, age, T stage, N stage, M stage, AJCC8 stage ^50^, BMI, smoker status, and alcohol use status. We used the R package gtsummary to perform Fisher’s exact tests for gender and self-reported race, and Kruskal-Wallis rank sum test for continuous or ordered categorical variables between HPVint(+) and HPVint(-).

### Identifying and visualizing the human genes with recurrent integration events

After annotating all integration events to genes and their flanking genes, we defined genes that appeared in at least two participants as “recurrent genes”. The genomic locations of integration events, along with their genomic annotations from above and which cohort they were from, were visualized using karyoploteR ^51^. We combined proximal (within a 1MB window) genes into “Gene clusters” for visualization.

Alex et al ^33^ collected recurrent HPVint hotspots in cervical cancer using a 3 MB window. Thus, we also collapsed all integration sites in our recurrent gene clusters into 3MB windows for comparability and overlapped our hotspots with theirs, using the Bioconductor package *GenomicRanges* with the function findoverlaps.

### Using STRING and BioGRID to identify protein-protein interactions

We used STRING ^52^ and BioGRID ^53^ to generate a protein-protein interaction (PPI) network with recurrent genes and integrated expression outlier genes. Visualization was performed using Cytoscape (v3.9.1). Genes were colored by the number of samples with integration present, shape was determined by expression outlier status, border color was used to annotate genes as known tumor suppressors or oncogenes or both based on the OncoKB ^54^, and edge color was determined by the database(s) supporting the PPI.

### HPV gene expression, E2:E6 ratio and ImmunOnco score calculation

We calculated HPV gene expression for each sample and correlations among HPV genes (See Supplementary Methods).

The ratio between E2 and E6 expression was calculated as:

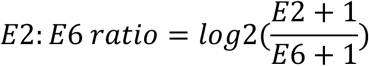

where *E2* and *E6* are the read counts. A value of 1 was added to avoid zeros. The ImmunOnco score was defined as:

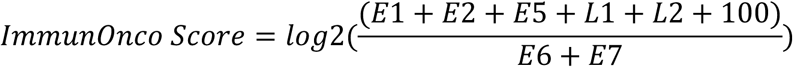

We added 100 to the numerator to mitigate the effect of extreme negative values after the log_2_ transformation. The Short ImmunOnco score was calculated as:

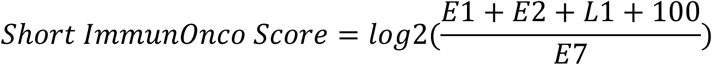

To eliminate batch effects among cohorts, we harmonized the ImmunOnco Scores. We first centered the individual scores (IO_i,c_) of each cohort by its mean (*μ*_c_), and then added the overall mean (*μ*_a_) back for the final score:

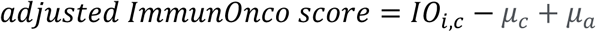

The ImmunOnco scores were calculated for 261 samples and 251 participants/cell lines. Sample TCGA-BB-7864-01A was infected by HPV68 and was removed for this analysis due to the lack of an official HPV68 gene annotation.

### Calculate the associations of cell type proportions with HPV integration status, E1* integration status, and ImmunOnco scores

Cell type proportions from bulk RNA-seq data were generated from CIBERSORTx (See Supplementary methods). Correlations between cell type proportions and HPV integration status, adjusted ImmunOnco scores (225 participants, TCGA-BB-7864-01A and the 10 duplicated samples were removed), and E1* integration status (130 HPVint(+) participants) were calculated. CD4+ and CD8+ T cell proportions were summed to define total T cells.

Histograms and percentages of zeros were analyzed for each cell type across samples to determine the appropriate transformation and analysis method. (**Supplementary Figure 2**, See Supplementary methods). Linear regression was used to test associations between continuous cell types and adjusted ImmunOnco scores, ANOVA was used for HPVint status and E1* integration status, and logistic regression was applied for cell types transformed to binary variables. All above analyses included cohort as a covariate. FDR-adjusted p-values were calculated using the Benjamin and Hochberg (BH) method and FDR < 0.05 was recognized as significant.

### Survival and recurrence analyses

We performed both log-rank test based on the Kaplan-Meier (KM) curve and multivariable Cox Proportional Hazards (CoxPH) regression to analyze overall survival and recurrence for HPV integration status, ImmunOnco score, and E1* integration status. The recurrence time is defined as the time to first recurrence or set to zero if the participant was never tumor-free (residual disease detected). We tested: 1) HPVint(+) with recurrent sites versus HPVint(+) no recurrent sites and HPVint(-). 2) HPVint(+) versus HPVint(-), 3a) HPV E1* integration versus HPVint(+) but not E1* integration and 3b) HPV E1* integration versus HPVint(-) plus HPVint(+) but not E1* integration , 4) ImmunOnco scores and its tertiles (standard and short version).

All ImmunOnco score-related analyses were performed for FF and FFPE separately, due to a batch effect between the FFPE and FF cohorts. All recurrence analyses focused on the 61 UM_FFPE patients since recurrence information was not available for the other cohorts. Multivariate CoxPH for recurrence included age and AJCC8 staging as covariates. We also tested the association of log2(E2/E6), log2(E1/(E6+E7)), log2((E2+E5)/(E6+E7)), log2((L1+L2)/(E6+E7)), E6 log2CPM, and E7 log2CPM values versus recurrence. For survival analyses, we utilized overall survival time and death status from 135 patients, comprising 66 TCGA, 18 UM_FF and 52 UM_FFPE participants (9 UM_FFPE overlapped with UM_FF). Multivariable CoxPH for survival incorporated age, AJCC8 staging, and smoker status as covariates. All KM curves and log-rank tests were conducted with the survfit2 function from ggsurvfit2 ^55^ (v0.3.0). The CoxPH regression analyses were performed using the coxph function from survival (v3.5.7)^56^.

### Comparison of sample-level HPV integration status by DNA versus RNA

Differences between samples positive for HPV integration by DNA versus RNA were assessed using the 102 participants (83 HVC and 19 TCGA) with both whole-genome sequencing (WGS) and RNA-seq data. DNA HPV integration events and status were from WGS data, as defined by Symer DE, et al^9^ but using our method for target gene assignments. Participants were categorized into four groups: double-positive (DNA+, RNA+); DNA-positive only (DNA+, RNA-); double-negative (DNA-, RNA-); and RNA-positive only (DNA-, RNA+). We also defined two types of HPV integration events, productive and silent (see Supplementary Methods).

### Differential expression and gene set enrichment analysis

To identify differentially expressed genes (DEGs) between double-positive (DNA+, RNA+) and double-negative (DNA-, RNA-) samples, we first adjusted raw gene counts from the 102 participants to remove batch effects between TCGA and HVC using Combat-seq. We used edgeR glmQLFTest with cohort as a covariate, and DEGs were defined as those with FDR<0.05.

For DEG analysis involving both FF and FFPE samples, we performed analyses stratified by cohort (UM_FF, UM_FFPE, TCGA, HVC) to avoid batch effects. We used the edgeR glmQLFTest for each cohort and we included “batch” as a covariate for the UM_FFPE four batches. We compared HPVint(+) versus HPVint(-), tertile high versus tertile low ImmunoOnco score tertiles, and E1* integration versus not E1* integration and HPVint(-). To identify consistent DEGs across the four cohorts (UM_FF, UM_FFPE, TCGA, and HVC), we used Fisher’s method to calculate meta p-values and then adjusted them using BH (FDR). Genes with FDR< 0.05 and greater than 50% change (logFC>0.58) in the same direction in at least three cohorts were displayed in the heatmaps.

For each comparison, we performed gene set enrichment testing using the enrichGO function from clusterProfiler 4.2.2 ^57^ with GOBP, GPMF, GOCC, and MsigDB Hallmark cancer gene sets based on genes with meta p-value < 0.05 and greater than 50% change in the same direction in at least three cohorts.

## Results

### Overview of HPV integration and cohort demographics

In this multi-cohort analysis, we analyzed 261 HPV(+) HNSCC RNA-seq samples from 249 participants and 3 cell lines, identifying a total of 1,117 high quality HPV-human fusion events. Each event was classified as unspliced or spliced, based on whether the HPV junction was at a common HPV splice site, resulting in 550 unique sample-human-location unspliced HPV insertion events and 567 spliced events (**Supplemental Table S1**). Each sample was classified as either HPVint(+), not determined (HPVint-ND), or HPVint(-) based on evidence for at least two confident human-HPV fusion reads (See Methods). The integration events among the 149 (57%) HPVint(+) samples were linked to 1,416 potentially affected unique human genes (**Supplementary Table S1**). Although 57% of samples were confirmed HPVint(+), 67% (HPVint(+) and HPVint-ND) had some evidence of integration, indicating additional samples having potential low-level expression of HPVint events (**Figure 1A, Supplementary Table S2**).

**Figure 1.**
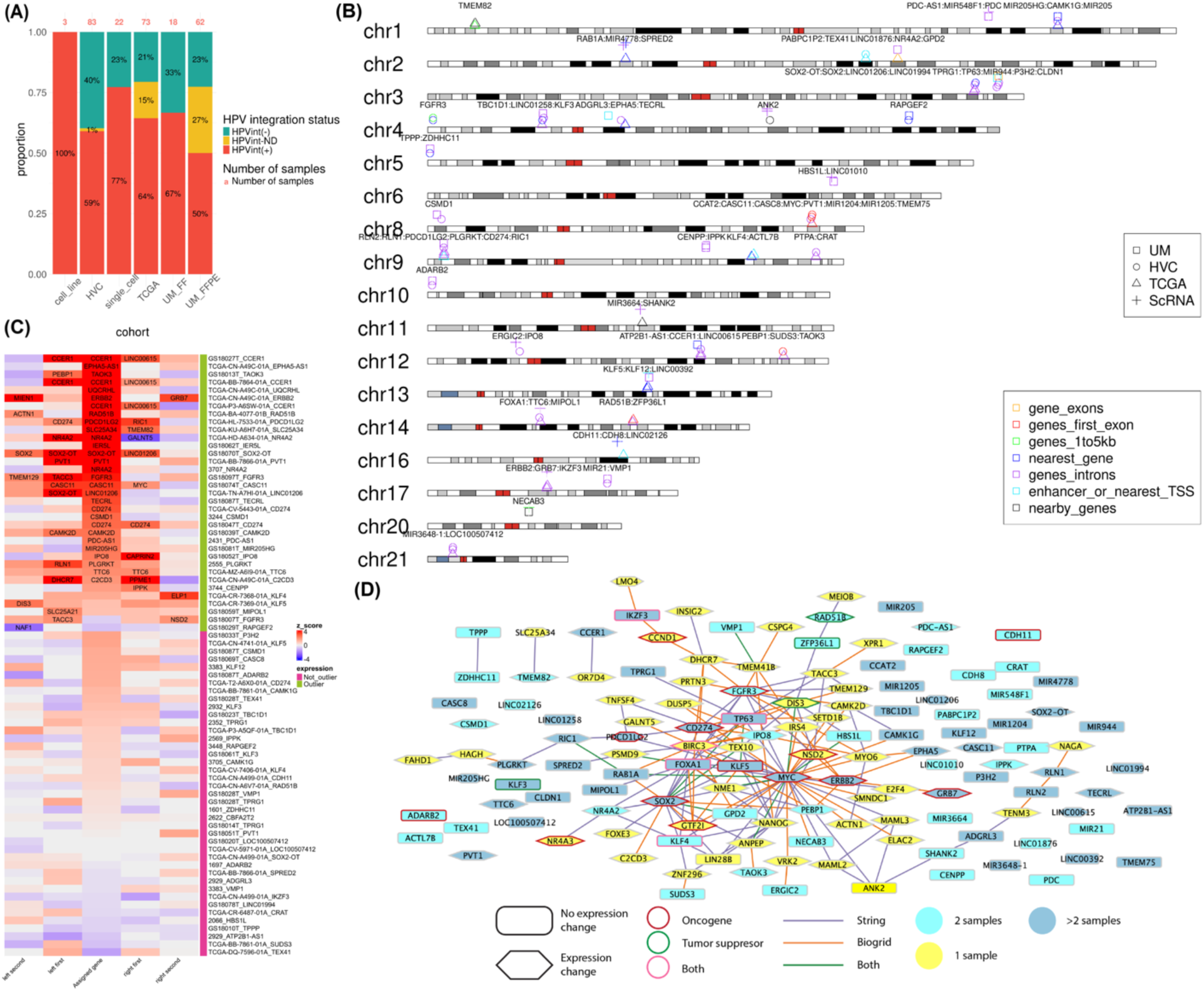
HPV integration landscape across HNSCC cohorts. (A) Stacked bar plot illustrating 6 analyzed HPV(+) HNSCC cohorts and their HPV integration status distribution. (B) Karyotype plot for all 34 HPV integration recurrent gene clusters. Different shapes represent cohorts and colors represent their gene annotations which are most relevant to expression. (C) Gene expression heatmap (Z-scores) for recurrent gene clusters. Each row represents a unique sample-gene pair. The middle column represents the ‘integrated genes’ (See Methods). The remaining columns are positional ‘nearby genes’. The heatmap is sorted by the middle column and annotated by the existence of expression outliers (|Z-score|>2). Gene symbols are noted for expression outliers. (D) Protein-protein interaction network among all recurrently integrated genes and integrated expression outlier genes. Genes are colored based on the number of participants, shape represents expression change status, border color annotates genes as known tumor suppressors or oncogenes, and edge color represents which database(s) supported the interaction.

Examining the potential associations between clinicodemographic information (**Supplementary Table S3**) and HPV integration status, we did not observe any statistically significant results, although HPVint(+) patients tended to have a lower tumor nodal (n) stage compared to HPVint(-) patients (Kruskal-Wallis rank sum test, *p*=0.063) (**Table 1**).

**Table 1.** Demographic information for all participants. Statistical tests were performed between HPVint(+) and HPVint(-)

| Variable | N | HPVint(+), N =<br>1451 | HPVint(-), N =<br>711 | p-value<sup>2</sup> |
| --- | --- | --- | --- | --- |
| gender | 213 |  |  | 0.11 |
| Female |  | 10 (7.0%) | 1 (1.4%) |  |
| Male |  | 133 (93%) | 69 (99%) |  |
| race | 112 |  |  | 0.8 |
| 1 |  | 75 (95%) | 32 (97%) |  |
| 2 |  | 2 (2.5%) | 0 (0%) |  |
| 5 |  | 1 (1.3%) | 1 (3.0%) |  |
| 6 |  | 1 (1.3%) | 0 (0%) |  |
| age | 131 | 58 (53, 66) | 58 (54, 60) | 0.6 |
| t_stage | 123 |  |  | 0.9 |
| 1 |  | 18 (21%) | 7 (19%) |  |
| 2 |  | 36 (41%) | 15 (42%) |  |
| 3 |  | 15 (17%) | 6 (17%) |  |
| 4 |  | 18 (21%) | 8 (22%) |  |
| n_stage | 124 |  |  | 0.063 |
| 0 |  | 16 (18%) | 3 (8.6%) |  |
| 1 |  | 16 (18%) | 5 (14%) |  |
| 2 |  | 52 (58%) | 22 (63%) |  |
| 3 |  | 5 (5.6%) | 5 (14%) |  |
| m_stage | 120 |  |  | 0.5 |
| 0 |  | 85 (99%) | 34 (100%) |  |
| 1 |  | 1 (1.2%) | 0 (0%) |  |
| AJCC8_stage | 109 |  |  | >0.9 |
| 1 |  | 27 (35%) | 13 (41%) |  |
| 2 |  | 30 (39%) | 9 (28%) |  |
| 3 |  | 20 (26%) | 9 (28%) |  |
| 4 |  | 0 (0%) | 1 (3.1%) |  |
| bmi | 51 | 27.1 (24.3, 31.7) | 29.8 (26.6, 33.6) | 0.3 |
| smoker | 129 |  |  | 0.9 |
| 0 |  | 41 (44%) | 15 (42%) |  |
| 1 |  | 9 (9.7%) | 4 (11%) |  |
| 2 |  | 43 (46%) | 17 (47%) |  |
| drinker | 119 |  |  | 0.5 |
| 0 |  | 19 (22%) | 5 (15%) |  |
| 1 |  | 23 (27%) | 10 (29%) |  |
| 2 |  | 43 (51%) | 19 (56%) |  |
| <sup>1</sup> n (%); Median (IQR) |  |  |  |  |
| <sup>2</sup> Fisher's exact test; Kruskal-Wallis rank sum test |  |  |  |  |

### Recurrent HPV integration in HNSCC pinpoints oncogenic networks and oncogenic expression

We used both unspliced and spliced integration events from the 149 HPVint(+) participants to generate a comprehensive HNSCC HPV integration map of recurrent genes (See Methods). Because insertion sites tend to cluster together in a tumor and may be associated with multiple nearby genes, we grouped those nearby genes into “gene clusters” based on their genomic locations (See Methods). Overall, 34 gene clusters (**Figure 1B, Supplementary Table S4)** were identified in at least two participants/cell lines and labeled as recurrent, with 14 appearing in three or more participants (**Table 2**). Seventy-five (50%) of the 149 HPVint(+) participants/cell lines were represented in a recurrent event, with 5 of them having two recurrent events and 5 having three or four. Among the 77 with clinical outcome information, there was no evidence for a difference in survival between patients with recurrent vs non-recurrent integration events (**Supplementary Figure 3A**). Of these 34 gene clusters, 31 (91%) were from at least two cohorts, suggesting they were not driven by batch or technical artifacts. Comparing our recurrent HPV integration gene clusters with 48 known cervical cancer recurrent “hotspots”,^33^ 17/34 (50%) overlapped (**Supplementary Figure 3B**), leaving 42 HNSCC unique recurrent related genes (in 17 recurrent gene clusters) not overlapping a CC recurrent gene cluster (**Supplementary Table S4)**. A closer examination reveals that the overlapping recurrent genes between CC and HNSCC include key oncogenic transcription factors (IKZF3, KLF5, MYC, SOX2, FOXA1, TP63), while the unique recurrent genes in HNSCC are associated with immune responses (CD274, PDCD1LG2) or beta-catenin binding (CDH8, CDH11, KLF4, NR4A2).

**Table 2.** Recurrent gene clusters appearing in at least three participants, bolded genes are annotated as tumor suppressors or oncogenes by OncoKB.

| Gene cluster | number of patients | patients | chr | start |
| --- | --- | --- | --- | --- |
| RLN2, RLN1, **PDCD1LG2**, PLGRKT, **CD274**, RIC1 | 5 | 2555,GS18047T,TCGA-CV-5443-01A,TCGA-HL-7533-01A,TCGA-T2-A6X0-01A | chr9 | 5399904 |
| TPRG1, **TP63**, MIR944, P3H2, CLDN1 | 5 | 130-MCDM-3,2352,GS18014T,GS18028T,GS18033T | chr3 | 189895060 |
| ATP2B1-AS1, CCER1, LINC00615 | 4 | 2929,GS18027T,TCGA-BB-7864-01A,TCGA-P3-A6SW-01A | chr12 | 89764269 |
| CCAT2, CASC11, CASC8, **MYC**, PVT1, MIR1204, MIR1205, TMEM75 | 4 | GS18051T,GS18069T,GS18074T,TCGA-BB-7866-01A | chr8 | 127854972 |
| **KLF5**, KLF12, LINC00392 | 4 | 3383,Sc_OP33,TCGA-CN-4741-01A,TCGA-CR-7369-01A | chr13 | 73731063 |
| SOX2-OT, **SOX2**, LINC01206, LINC01994 | 4 | GS18070T,GS18078T,TCGA-CN-A499-01A,TCGA-TN-A7HI-01A | chr3 | 181989048 |
| TBC1D1, LINC01258, **KLF3** | 4 | 2932,GS18023T,GS18061T,TCGA-P3-A5QF-01A | chr4 | 38431916 |
| ADGRL3, EPHA5, TECRL | 3 | 2929,GS18087T,TCGA-CN-A49C-01A | chr4 | 60105050 |
| ANK2 | 3 | GS18039T,Sc_OP17,Sc_OP35 | chr4 | 113837077 |
| **ERBB2, GRB7,**<br>**IKZF3** | 3 | Sc_GEX_HN68,TCGA-CN-A499-01A,TCGA-CN-A49C-01A | chr17 | 39742251 |
| **FOXA1**, TTC6,<br>MIPOL1 | 3 | GS18059T,Sc_C22,TCGA-MZ-A6I9-01A | chr14 | 37238783 |
| MIR205HG,<br>CAMK1G,<br>MIR205 | 3 | 3705,GS18081T,TCGA-BB-7861-01A | chr1 | 209519796 |
| MIR3648-1,<br>LOC100507412 | 3 | GS18001T,GS18020T,TCGA-CV-5971-01A | chr21 | 8438760 |
| RAB1A,<br>MIR4778,<br>SPRED2 | 3 | Sc_HN66,Sc_OP6,TCGA-BB-7866-01A | chr2 | 65906915 |

To assess the impact of HPV integration events on the expression of surrounding genes, we first compared the expression of the gene assigned to each integration event (termed ‘integrated gene’, see Methods) in the integrated sample(s) with all other samples. For this, we used Z-scores stratified by cohort for all bulk RNA-seq HPVint(+) participants. We found that 46% of the integrated genes had outlier expression levels (absolute value of Z-score>2) for both recurrent and non-recurrent genes, (**Figure 1C, Supplementary Figure 3C**), and the great majority (167/226; 74%) of integrated genes had above average expression (Z-score>0) (**Figure 1C, Supplementary Figure 3C**). Within recurrent gene clusters, well-known oncogenes NR4A2, CD274, and CCER1, appeared multiple times across different patients as expression outliers (NR4A2: 2 of 2 times, CD274: 3 of 5 times, CCER1: 3 of 3 times), indicating their importance as oncogenes in HPV(+) HNSCC (**Figure 1C**). Recurrent oncogenes from the CAMK (CAMK2D, CAMK1G) which are unique to HNSCC and KLF (KLF3, KLF4, KLF5, and KLF12), of which KLF3 and KLF4 overlap with CC, families also tended to be upregulated due to HPV integration (one sample t-test for Z-scores>0, p=1.04e^−7^). Together, these results confirm that HPV integration in HNSCC directly contributes to the overexpression of human oncogenes and is involved in carcinogenesis, with substantial similarity with CC (**Figure 1C**).

To further explore the genes and pathways impacted by HPV integration, we built a protein-protein interaction (PPI) network showing interactions among all recurrent genes and any integrated expression outlier gene (absolute value of Z-score>2) having an interaction with the others. Recurrent genes CD274, BIRC3, FOXA1, SOX2, GTF2I, NME1, KLF5, TEX10, TP63, FGFR3, IPO8, DIS3, IRS4, NSD2, MYC, NANOG, ERBB2,

TMEM129 and ANK2 are highly connected to the center hub of the network indicating their importance (**Figure 1D**). Interestingly, the genes ERBB2 and NR4A2 have not been previously reported as recurrent genes but were identified as frequently mutated genes in cancers^8^. In addition, we identified 14 expression outliers (but not recurrent genes) with more than 3 protein interactions, all but one of which are recognized as an oncogene in one or more cancer types (**Supplementary Table S4**). Our PPI summarizes a comprehensive, HNSCC-specific HPV integration-related oncogenic network, offering a valuable reference for future targeted therapy.

### Classification of HPV integration events based on splicing and HPV gene ratios

HPV integration status has been suggested as a potential biomarker for risk stratification in HPV+ OPSCC, yet the literature is mixed regarding its relationship with clinical outcomes, suggesting more accurate stratification is needed^7^. HPV integration events often generate human-HPV spliced mature mRNA ^9,38^, and our meta-cohort shows frequent splicing at or near the HPV16 genome position 881 which is located at gene E1 (**Figure 2A-B**). We categorized samples having E1-related spliced site events as having ‘E1* integration’ (110 of 151 (73%) samples within +/-2 bp), and similarly classified samples with an E6*-related integration event (position 227 for HPV16) as having ‘E6* integration’ (51 (34%) samples within +/-2 bp) (See Supplementary Methods). We observed a strong correlation between E1* integration and E6* integration participants, with E1* integration appearing to act as a gatekeeper to E6* integration (Fisher exact test: OR=6.83, p-value=8.58e^−05^) (**Table 3**). Upon closer inspection of the four exception cases to this rule (i.e. E6* integration without E1* integration), we found 2 of the 4 had potential E1* spliced integration events that were filtered out by the integration software, suggesting at least two of these were likely false negatives. The 110 E1* integration participants had significantly lower E2:E6 compared to other participants overall (logistic regression p-value=7.45e^−8^ with cohort as a covariate) and for 3 cohorts individually (all except for UM_FF, which had the smallest sample size, and TCGA which had shorter single-end reads; both UM_FF and TCGA demonstrated the same trend) (**Figure 2C**). No significant E2:E6 difference was observed between HPVint(+) participants without E1* integration and HPVint(-) participants, and E2:E6 was not as strongly associated with E6* integration (**Figure 2C-D**). Additionally, at the single gene level of all 1416 HPV integration related genes in this study, E1* integration was highly associated with human recurrent integration genes (Fisher exact test: OR=6.29, p-value=6.77e^−13^). These results indicate that i) HPVint can generate spliced HPV-human fusion transcripts spliced at specific HPV sites, ii) that splicing of E1* likely occurs before splicing of E6* and is associated with loss of E2, and iii) that these spliced integration transcripts are more likely to happen when integration occurs at a “hotspot” site.

**Figure 2.**
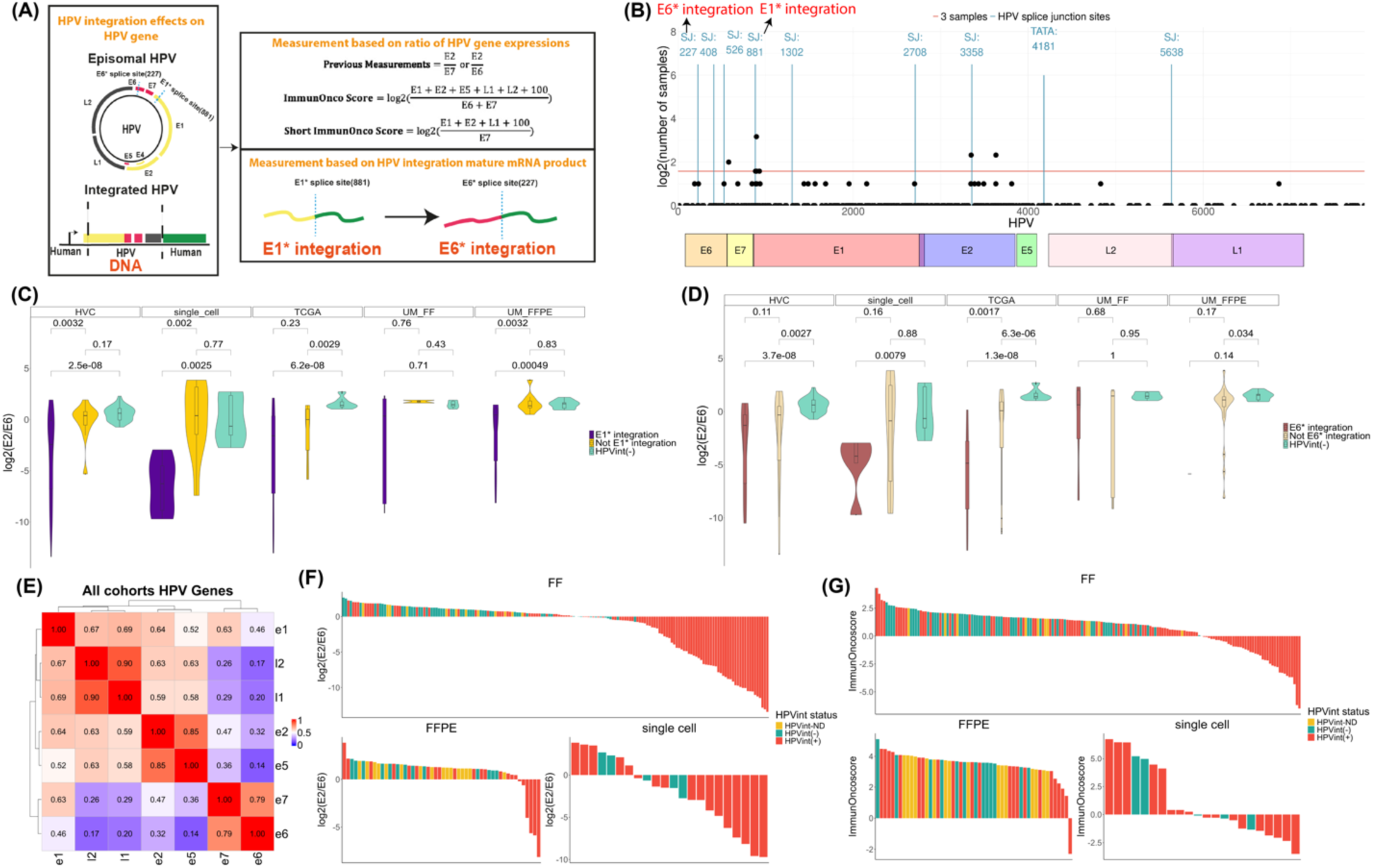
Novel strategies to classify HPV integration (+) HNSCC participants. (A) Schematic describing new classification strategies for HPV integration status by i) HPV gene expression ratios and ii) mature spliced mRNA products. (B) RNA-based HPV integration junctions across the HPV genome, showing enrichment at HPV16 known splice junctions. (C) Violin plots across cohorts showing the log(E2/E6) among E1* integration, Not E1* integration, and HPVint(-) groups. (D) Violin plots across cohorts showing the log_2_(E2/E6) among E6* integration, Not E6* integration and HPVint(-) groups. (E) Correlation heatmap among HPV genes using log_2_cpm values for all samples. (F)-(G) Waterfall plots illustrating log_2_(E2/E6) and ImmunOnco score, colored by HPV integration status and separated by tissue/sequencing type (fresh frozen (FF), FFPE, or single cell).

**Table 3.** Fisher’s exact test for E1* integration versus E6* integration.

|  | E1* integration | Not E1* integration |
| --- | --- | --- |
| E6* integration | 47 | 4 |
| Not E6* integration | 66 | 62 |

Previous studies have tested the E2:E6 or E2:E7 ratio as a prognostic biomarker ^39,40^, however, we noticed that the expression of other HPV genes (E1, E5, L1, and L2) can also be lost to varying degrees irrespective of E2 loss. Thus, we explored new biomarker approaches by considering ratios involving more than E2 (**Figure 2A**). We hypothesized that expression loss of these relatively non-oncogenic HPV genes provides fewer antibody targets for the host immune system, leading to a reduction in tumor-infiltrating immune cells and more likely recurrence over time. Therefore, we defined a new score called the ImmunOnco score, based on the relative expression of relatively non-oncogenic (potentially immunogenic) to oncogenic genes, E6 and E7. We further examined the correlations among HPV genes and found that the expression of the pairs E2-E5, L1-L2, and E6-E7 were consistently highly correlated overall and among cohorts (log2CPM values, Pearson’s R values E2-E5:0.85, L1-L2:0.90, E6-E7:0.79; **Figure 2E, Supplementary Figure 4A**). Therefore, we also created a simplified score, using E1, E2, L1, and E6 only called the Short ImmunOnco score (**Supplementary Table S5**). As expected, both lower E2:E6 and lower ImmunOnco score were indicators for the occurrence of HPV integration (**Figure 2F-G**). E1* integration was also associated with lower ImmunOnco score in all cohorts (significant in TCGA *p*=0.045, HVC: *p*=0.012, and single cell *p*=0.0068), however, E6* integration was only significant for TCGA and single-cell cohorts and with inconsistent trends among cohorts (**Supplementary Figure 4B-C**).

### E1* integration is associated with worse survival while ImmunOnco score is associated with recurrence

To investigate whether clinical outcomes differ among patients with different consequences of HPV integration, we collected participants’ overall survival (n=135) and recurrence information (n=70) (**Supplementary Table S5**) and performed both log-rank tests with Kaplan-Meier curves and multivariable Cox Proportional Hazards (CoxPH) regression analyses (see Methods).

For overall survival based on HPVint status, we observed a non-significant trend for HPVint(+) participants having worse survival than HPVint(-) participants using both the log-rank test (p=0.095) (**Supplementary Figure 5A**) and multivariable CoxPH analysis (HR=3.724, p=0.079) (**Supplementary Figure 5B**). E1* integration participants, however, had significantly worse survival compared to HPVint(-) participants or all other HPV(+) patients (HPVint(-) and Not E1* integration) (log-rank test *p*=0.002, HR=4.977 & 8.094, CoxPH *p*=0.032 & 0.0054, respectively) (**Figure 3A-B, Supplementary Figure 5C**), suggesting that E1* integration is a potential biomarker for poor prognosis and clinical risk stratification in HPV+ OPSCC.

**Figure 3.**
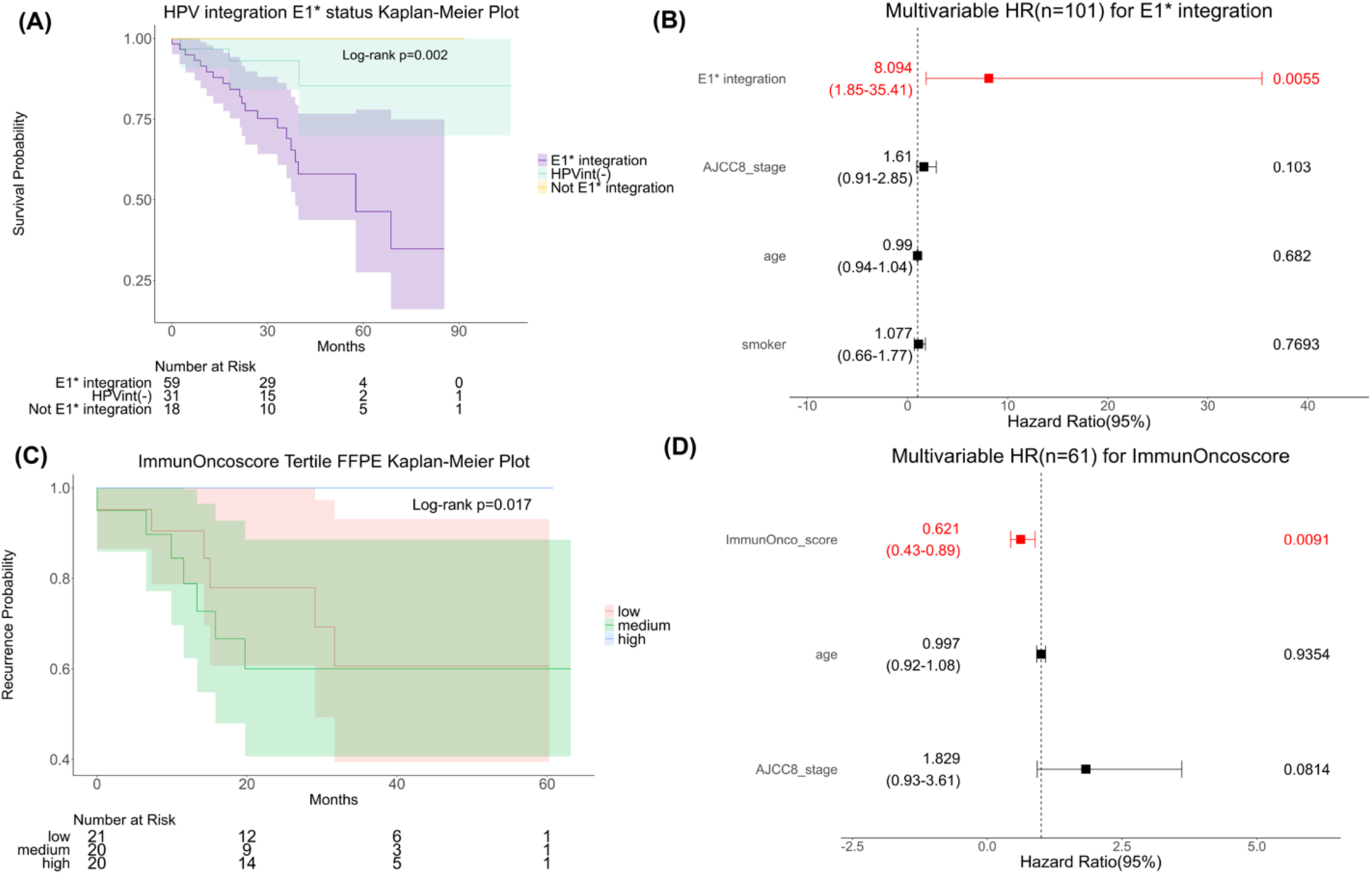
Clinical outcomes are associated with E1* integration and ImmunOnco score. (A) Overall survival Kaplan-Meier (KM) plot for E1* integration, Not E1* integration and HPVint(-), with log-rank p-value across three groups. (B) Overall survival Forest plot describing multivariable CoxPH regression for E1* integration versus Not E1* integration including HPVint(-) status, with AJCC v8 stage, age, and smoking status as covariates. (C) Tumor recurrence KM plot for ImmunOnco score, with log-rank p-value across tertiles. (D) Recurrence Forest plot describing multivariable CoxPH regression for ImmunOnco score, with AJCC v8 stage and age as covariates.

Due to a batch effect between fresh frozen and FFPE samples for the ImmunOnco score, we tested its association with survival stratified by this variable. For both 61 FFPE-based and 74 FF-based participants, ImmunOnco score was not significantly associated with survival (log-rank *p*=0.5 and 0.7; HR=0.885 and 1.009, CoxPH *p*=0.7007 and 0.9459, respectively) (**Supplementary Figure 5D-G**). However, the ImmunOnco score was significantly associated with higher tumor recurrence rate using a CoxPH model accounting for tumor stage (HR=0.621, *p*=0.00906) and the log-rank test (*p*=0.017) (**Figure 3C-D**). Similar results were found for the short ImmunOnco score (**Supplementary Figure 6A-B**) and for relative loss of other HPV genes (**Supplementary Figure 6C-E**). Recurrence significance with a CoxPH model did not extend to E2:E6 (HR=0.931, CoxPH *p*=0.32) (**Supplementary Figure 6F**) or to E1* integration (HR=0.694, CoxPH *p*=0.57), and was not due to higher E6 or E7 alone (**Supplementary Figure 6 G-H,** HR=1.112 and 1.093 and CoxPH *p*=0.63 and 0.73, respectively). Overall, these results demonstrate that the loss of non-oncogenic HPV gene expression (not just E2) and the biomarker E1* integration are predictive of worse clinical outcomes, occurrence of recurrence and worse survival, respectively.

### Keratinization, gated channel activities, immune response and infiltration are associated with downstream consequences of HPV integration

We conducted differential expression analyses and cell type deconvolution to investigate the genes, pathways, and immune cell types associated with E1* integration and high versus low ImmunOnco score (See Methods). We identified mostly distinct sets of DEGs for HPVint(+) versus HPVint(-) (466 genes); high versus low ImmunOnco score (259 genes), and E1* integration versus non-E1* integration and HPVint(-) (293 genes) (**Supplementary Table S6**). The differentially expressed genes identified from different comparisons follow similar trends across cohorts (**Figure 4A**). Over-represented pathways analysis identified three comparisons shared an enrichment of higher keratinization, cornified envelop and gap junction genes (**Figure 4B, Supplementary Table S7**). B cell proliferation and activation of immune response were uniquely upregulated in tumors with higher ImmunOnco score. (**Supplementary Table S7**). Downregulation of T cell receptor complex was most strikingly enriched with E1* integration (**Figure 4B, Supplementary Table S7)**. The results partially align with previous findings that HPV integration is associated with keratinization and immune response ^8^ and confirm that higher keratinization and cell junction changes are broadly associated with HPVint, but deactivation of the immune response is characteristic only of the subset of HPVint tumors with loss of non-oncogenic HPV gene expression, and lower T cell receptor complex expression is a signature of the subset with E1* integration.

**Figure 4.**
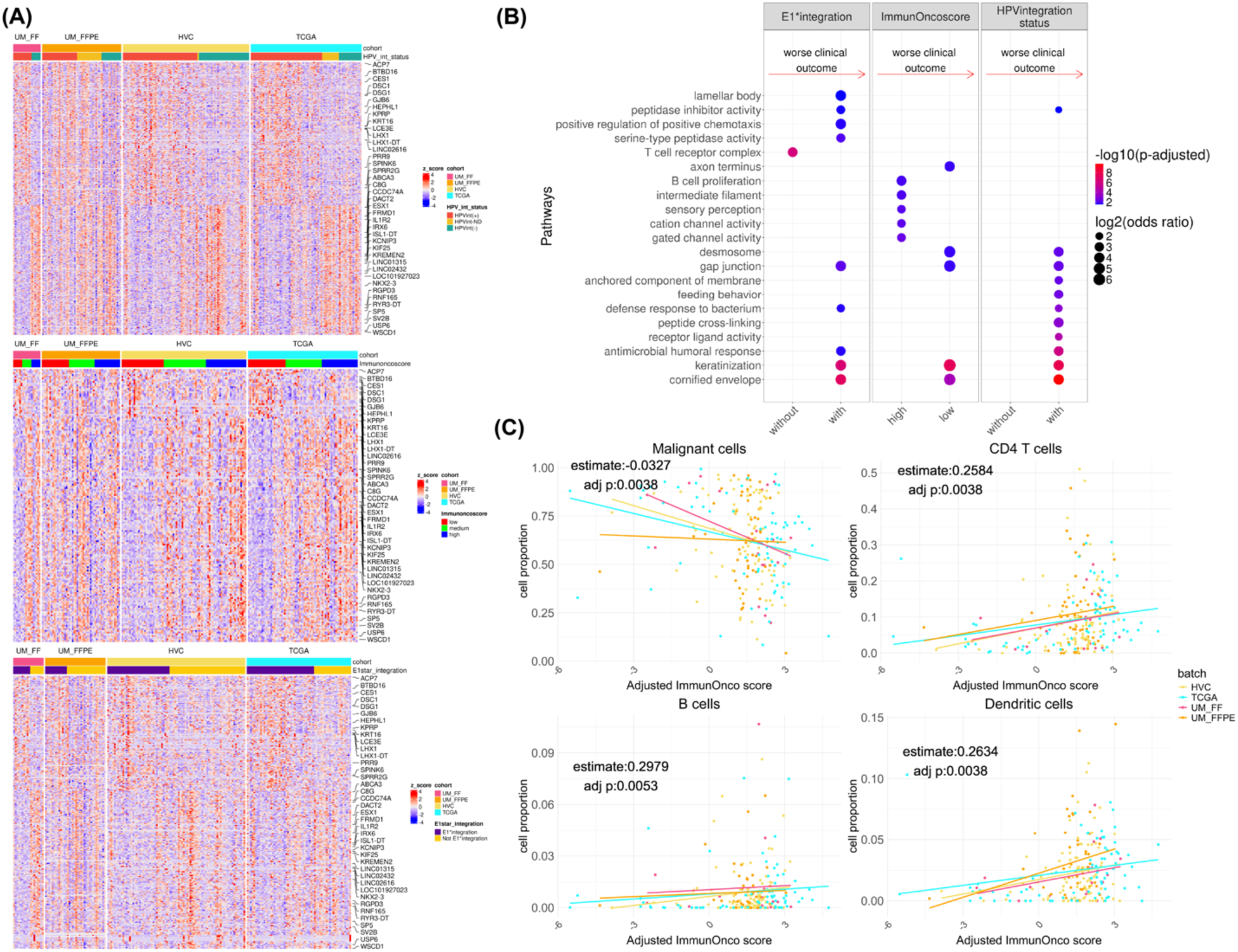
Differentially gene expression patterns and cell types by HPVint status, E1* integration, and ImmunOnco score. (A) Heatmaps of top differentially expressed genes (Z-scores; Fisher’s adjusted *p* <0.05, and at least 3 cohorts with the same direction abs(log_2_ fold change)>0.58). Top: HPVint(+) versus HPVint(-), middle: ImmunOnco score tertile high versus tertile low, bottom: E1* integration versus Not E1* integration and HPVint(-), genes annotated are shared among three comparisons (B) Gene set enrichment testing results from the above comparisons, top 10 significant GO terms ranked based on FDR are shown. (C) Scatterplots of adjusted ImmunOnco score versus four cell type proportions (Malignant cells, CD4+ T cells, B cells and Dendritic cells), colored by cohort.

To further examine the relationship between these downstream effects of HPVint and the tumor immune microenvironment, we tested association of the cell type composition with HPV integration status, ImmunOnco score, and E1* integration. Neither HPVint status nor E1* integration was significantly associated (FDR<0.05) with any of the ten cell types tested (**Supplementary Table S8**). ImmunOnco scores, however, were significantly positively associated with CD4+ T cells (adj p=0.0038), total T cells (adj p=0.0038), dendritic cells (adj p=0.0038), B cells (adj p=0.0052) and endothelial cells (adj p=0.014), and negatively associated with malignant cells (adj p =0.0038) (**Supplementary Table S8**), with the trend being consistent across cohorts (**Figure 4C**). These results further support our hypothesis that loss of non-oncogenic HPV gene expression as observed by a lower ImmunOnco score leads to a reduction in anti-tumoral immune infiltration.

### Concordance and discrepancies using DNA versus RNA for HPV integration detection

Previous studies of HPV integration in HNSCC are based on either DNA or RNA alone, potentially leading to discrepant reports regarding the downstream effects of HPV integration. Thus, we performed a comparison of the 102 samples (83 HVC and 19 TCGA) having both DNA (WGS) and RNA (RNA-seq) HPV integration detection. Among these samples, HPV integration status was consistent between RNA and DNA for 80 participants (79%), with 60 (59%) double-positive and 20 (20%) double-negative (See methods). Eighteen (18%) of the samples had exclusively DNA HPV integration, while only four (4%) had integration detected solely in RNA **(Figure 5A)**; of these four, two had confident HPVint spliced sites, likely indicating false negatives from the DNA.

**Figure 5.**
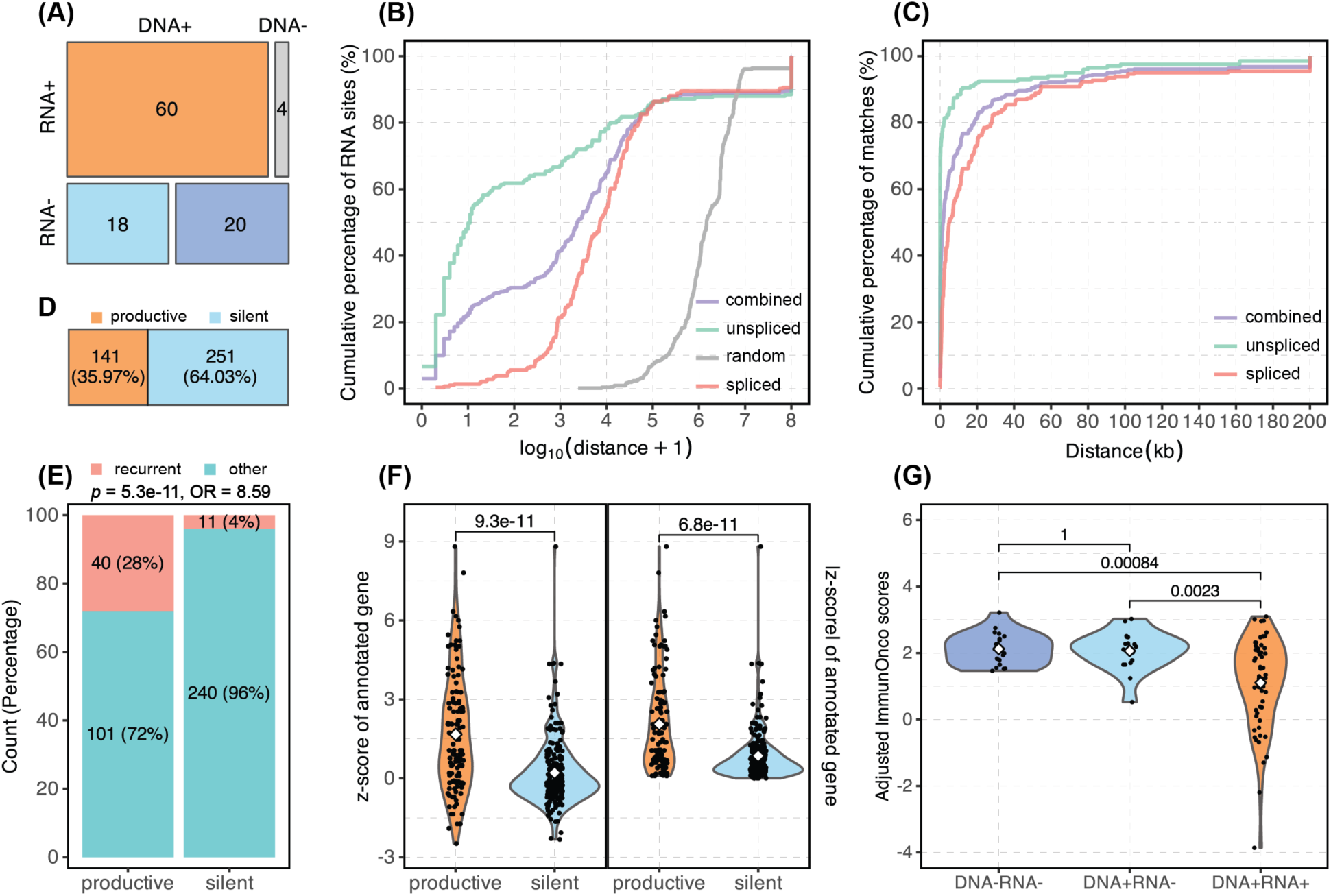
Characteristics of HPV integration events identified by DNA versus RNA. (**A**) Classification of patients based on HPV integration detection method. Color indicates patient group (DNA+RNA+: integration detected by both RNA-seq and WGS, DNA-RNA+: RNA-seq only, DNA+RNA-: WGS only, and DNA-RNA-: negative for both). (**B**) log-scale cumulative distribution plot of RNA fusion breakpoints illustrating the distances between RNA breakpoints and their proximal DNA breakpoints within samples. Combined is unspliced and spliced, random sites are shown in grey. (**C**) Cumulative concordance rate plot of RNA fusion breakpoints showing how well the most confidently annotated genes are matched by distance between RNA breakpoints and their nearest DNA breakpoints within samples. (**D**) Proportions of virus-host DNA breakpoints classified as productive (orange) and silent (light blue), based on the presence of RNA breakpoints within a ±100kb range of the DNA breakpoints. (**E**) Bar plot showing productive integration sites are significantly more likely to be annotated to a recurrent gene than silent integration sites. The *p*-value and odds ratio are for the two-tailed Fisher’s Exact test. (**F**) Violin plots illustrating the expression levels of the most confidently annotated genes at productive versus silent DNA breakpoints. Left plot shows the Z-score, right plot shows the absolute Z-score (|Z-score|). *P*-values are for the Student’s t-test. (**G**) Comparison of ImmunOnco scores among different patient integration categories (DNA-RNA-, DNA+RNA-, and DNA+DNA+), illustrated using violin plots. The *p*-values are for the Student’s t-test.

We examined the integration breakpoints identified by both DNA and RNA, detecting 739 DNA breakpoints and 511 RNA breakpoints among the 102 participants, including 225 (44%) unspliced RNA breakpoints and 286 (56%) spliced RNA breakpoints. Approximately 10% of RNA breakpoints (53 = 26 unspliced +27 spliced) failed to map to a DNA breakpoint within 100 Mb on the same chromosome. As expected, unspliced RNA breakpoints were much closer to a DNA breakpoint (median of 6 bp) than spliced RNA breakpoints (median of 4,846 bp). Overall, ∼85% of the RNA breakpoints were within 100 kb from a DNA breakpoint, which is much closer than observed for random genomic locations (**Figure 5B**). We observed >90% concordance in annotated genes between DNA and RNA breakpoints within a 60 kb range, narrowing to 15 kb for unspliced RNA breakpoints (**Figure 5C**). This high concordance rate supports the inclusion of spliced RNA integration events and indicates consistency between DNA and RNA HPV integration for gene-level analysis.

After merging breakpoints that were closer than 10 kb and annotated to the same gene, 392 DNA integration sites remained. About 64% of these were classified as silent (not observed in RNA), while 36% were productive (observed in RNA) (**Figure 5D**, See methods). Notably, 40 of the 141 productive sites (28%) corresponded to recurrent genes defined above, which was seven times higher than that observed for silent sites (4%), indicating a strong selective pressure for recurrent genes to produce fusion RNAs (*p* = 5.28 × 10^−11^, OR = 8.59, two-tailed Fisher’s exact test) (**Figure 5E**). Moreover, expression levels of the annotated gene at productive sites were significantly different from those at silent sites, with genes at productive sites showing increased expression (Z-score, *p* = 9.3 × 10^−11^; |Z-score|, *p* = 6.8 × 10^−11^, Student’s t-test) whereas the median Z-score for silent site genes was 0.21 (median |Z-score| for silent site genes was 0.57) indicating minimal shifts in expression (**Figure 5F**). One may hypothesize that silent events occur more often in exons or 5’UTRs, disrupting a tumor suppressor gene. However, we found 15 (11%) of productive sites and only 10 (4%) of silent sites in an exon or 5’UTR. Consequently, we did not find any evidence that silent sites were enriched in regions likely to disrupt gene function. Together, these findings underscore the more limited relevance of integration events detected solely in DNA to carcinogenesis.

To investigate host gene expression differences by HPVint status based on DNA and RNA, we first tested host differential expression between double-positive and double-negative tumors and identified 220 DEGs (FDR < 0.05). Hierarchical clustering showed DNA-positive only tumors were almost evenly split between clustering with double-positive and double-negative tumors (**Supplementary Figure 7A**), further supported by principal component analysis (PCA) (**Supplementary Figure 7B**). However, assessing the expression of HPV genes, the adjusted ImmunOnco scores were significantly higher in double-negative (*p* = 8.4 × 10^−4^, Student’s t-test) and DNA-positive only patients (*p* = 2.3 × 10^−3^, Student’s t-test) than in double-positive patients, while no difference was observed between DNA-positive only and double-negative patients (*p* = 1, Student’s t-test) (**Figure 5G**). This result confirms that RNA expression from the integration event is needed for loss of non-oncogenic HPV genes.

## Discussion

Human papillomavirus DNA integrated into the human genome often occurs early in tumor development but continues through progression ^58^. Many studies in cervical cancer ^35,59^ and HNSCC ^9,12^ have elucidated the DNA-level consequences of HPV integration. However, the downstream impact of integration on the host and HPV gene expression levels and splicing, and how these affect clinical outcomes, remained unclear. We analyzed RNA-seq data from 261 HPV(+) HNSCC samples, assessing the gene expression, splicing, and clinical impacts of RNA level integration, finding evidence to explain previous discrepancies in literature. Further, we expanded our cohorts by introducing a customized approach to detect viral-host fusion transcripts in single cell RNA-seq data.

In doing so, we also provide the most comprehensive resource of genes with recurrent HPV integration events in HNSCC. Recurrent HPV integration sites have been well summarized in cervical cancer ^33^ but not in HNSCC due to insufficient data. Compared to CC, we discovered 17 new recurrent gene clusters. Our protein interaction network with recurrent and integrated expression outlier genes captured previously mentioned essential onco- or tumor suppressor genes: SOX2, KLF5,FOXA1, KLF5, MYC, BIRC3,TP63, and FGFR3 ^8,9^, but also highlighted novel ones including DIS3, NANOG, NME1, NSD2, and ERBB2. Most of the genes in protein interaction network are transcription factors and highly interact with each other, indicating a single disturbance may perturb an entire oncogenic network. By comparing with previous DNA-based HPint results ^9^, we found that integration events linked to recurrent genes are 7-times more likely to express human-HPV fusion RNA than non-recurrent genes. These findings highlight the importance of summarizing these recurrences at the RNA level to evaluate potential HPV integration consequences.

Previous studies mentioned but did not systematically evaluate the influence of HPV integration on HPV gene splicing ^9,38,60^ . Our study evaluated these HPV features to identify potential biomarkers associated with worse clinical outcomes in subsets of HPVint(+) participants. Building on the knowledge that E2:E6 ^39^ and E2:E7 ^61^ are indicators of worse prognosis, and E2 expression is regulated by site 881 splicing in HPV16 as an example^20^, we further classified patients with HPV integration based on whether spliced integration events were detected near 881 (E1* integration), and an extended score of E2:E6 using all HPV non-oncogenic genes (ImmunOnco score). This extension is driven by the hypothesis that all HPV genes could be seropositive ^62,63^ and E6 seropositivity has been linked not only to survival ^63^ but also to recurrence ^64^. We discovered E1* integration is associated with E2:E6 ratio and acts as a gatekeeper for E6* integration, and hypothesized that it may derive from spliced mature human-HPV mRNA. More importantly, our classifications partially address the conflicting results regarding the association between HPV integration and clinical outcomes. Some previous studies found an insignificant trend towards worse survival in patients with HPV integration ^2,9^. Our results suggest that it is only a subset of HPVint(+) patients who have worse prognosis, i.e. those with E1* splicing integration events.

Previous studies also overlooked the potential impact of HPV gene expression ratios on clinical outcomes. We expanded the E2:E6 or E2:E7 ratio to more broadly consider relatively non-oncogenic to oncogenic gene ratios. For this work, we classified E5 as a potentially immunogenic gene as opposed to oncogenic, which may be controversial since E5 plays a supportive role in carcinogenesis by cooperating with E6 and E7 and promoting immune evasion and proliferation ^65^. However, little is known regarding the consequences of E5 loss upon HPV integration and given the strong correlation in expression between E2 and E5 across tumors, we could not effectively distinguish their effects based on the RNA-seq data. Our findings indicate that participants who lost expression of any non-oncogenic (potentially immunogenic) gene (i.e. lower ImmunOnco score) were more likely to have tumor recurrence. We also demonstrated that the above two markers are correlated, with E1* integration tumors also having lower ImmunOnco scores. We were unable to answer the question of whether RNA HPVint is more predictive of worse prognosis than DNA HPVint, due to not having clinical outcome information for most patients with both DNA and RNA HPVint data, however Symer et al reported no association between DNA HPVint status and survival in the HVC cohort ^9^.

Leveraging our multiple cohorts, we found several key cancer pathways associated with HPV integration, ImmunOnco score, or E1* integration. Keratinization was heightened in both HPVint(+) and low ImmunOnco score participants, which is also a characteristic of HPV(-) HNSCC more broadly. Low ImmunOnco scores were associated with downregulation of lymph node development, immune response activation, and fewer infiltrating B cells, T cells, and dendritic cells, providing evidence for the hypothesis that continued expression of non-oncogenic HPV genes leads to a more immunogenic tumor, more likely classified as an immune ‘hot’ phenotype. E1* integration was uniquely associated with downregulation of T cell receptor complex and channel activities, potentially explaining the worse overall survival of participants with E1* integration.

Most HPV integration studies have focused on DNA events, yet our findings point to RNA as key to understanding how HPVint is associated with clinical outcomes. On one hand, splicing of integrated HPV RNA induces extrinsic RNA products such as E1* integration. On the other hand, based on our comparisons, DNA-only HPV integration may not significantly contribute to disturbance of HPV gene expression and is less likely to occur at recurrent sites. Unfortunately, the lack of available clinical outcome information for most of these participants precluded us from testing survival or recurrence differences between our defined double-positive, DNA-positive and double-negative participants.

Limitations of our study included heterogeneous tumor sample collection, limited clinical outcome availability, and a lack of diversity in terms of racial representation. The mix of fresh frozen and FFPE samples for RNA sequencing caused a batch effect in the ImmunOnco scores that could not be adequately corrected for using any simple transformation. This suggests that E1* integration may be more robust to differences in technical variations, and thus a more feasible clinical biomarker for survival. On the other hand, ImmunOnco score also motivates additional studies in larger cohorts, since as a continuous variable it may provide more power and flexibility to define meaningful clinical subgroups for precision oncology. ImmunOnco score was associated with recurrence, but recurrence information was only available for 70 of our 249 participants thus limiting our power to identify associations with it. In terms of cohort diversity, all but two of the UM cohort participants were non-Hispanic white, and we do not know the racial makeup of the HVC cohort. Future studies are needed to understand associations between race/ethnicity and the biomarkers examined in this manuscript.

Our study comprehensively illustrated the aberrant downstream consequences of HPV integration for both human and HPV RNA. We identified novel potential oncogenes in HPV-associated HNSCC through HPVint recurrence analysis and discovered potential clinical biomarkers, providing guidance for future HPV integration clinical-related studies. The ImmunOnco score and E1* integration were introduced as predictors of poor outcome. Yet future studies in larger cohorts are necessary to further assess these as prognostic biomarkers potentially translatable for precision oncology trials.

## Acknowledgements

This work was supported by National Institutes of Health grants R01-CA250214, T32-CA140044, and P01-CA240239. We acknowledge the Advanced Genomics Core at the University of Michigan. We also acknowledge the Ruben, et al. who shared their HPV (+) HNSCC single cell raw data. Additionally, this study makes use of data generated by Drs. Gillison, Symer and Akagi in the HPV Virome Consortium, formerly at the Ohio State University Comprehensive Cancer Center and now at University of Texas MD Anderson Cancer Center. Funding and computational support for these data were provided by the Ohio State University Comprehensive Cancer Center, the Ohio Supercomputer Center, Cancer Prevention & Research Institute of Texas and the University of Texas MD Anderson Cancer Center.

## Notes

Conflict of Interest Statement: The authors declare no potential conflicts of interest.

### Competing Interest Statement

The authors have declared no competing interest.

## References

1. Lechner, M., Liu, J., Masterson, L. & Fenton, T. R. HPV-associated oropharyngeal cancer: epidemiology, molecular biology and clinical management. Nat. Rev. Clin. Oncol. 19, 306–327 (2022).

2. Nulton, T. J., Kim, N. K., DiNardo, L. J., Morgan, I. M. & Windle, B. Patients with integrated HPV16 in head and neck cancer show poor survival. Oral Oncol. 80, (2018).

3. Menezes, F. D. S., Fernandes, G. A., Antunes, J. L. F., Villa, L. L. & Toporcov, T. N. Global incidence trends in head and neck cancer for HPV-related and -unrelated subsites: A systematic review of population-based studies. Oral Oncol. 115, 105177 (2021).

4. Economopoulou, P., Kotsantis, I. & Psyrri, A. De-Escalating Strategies in HPV-Associated Head and Neck Squamous Cell Carcinoma. Viruses 13, (2021).

5. McBride, A. A. & Warburton, A. The role of integration in oncogenic progression of HPV-associated cancers. PLoS Pathog. 13, e1006211 (2017).

6. Karimzadeh, M. et al. Human papillomavirus integration transforms chromatin to drive oncogenesis. Genome Biol. 24, 142 (2023).

7. Tabatabaeian, H., Bai, Y., Huang, R., Chaurasia, A. & Darido, C. Navigating therapeutic strategies: HPV classification in head and neck cancer. Br. J. Cancer (2024) doi:10.1038/s41416-024-02655-1.

8. Koneva, L. A. et al. HPV Integration in HNSCC Correlates with Survival Outcomes, Immune Response Signatures, and Candidate Drivers. Mol. Cancer Res. 16, 90–102 (2018).

9. Symer, D. E. et al. Diverse tumorigenic consequences of human papillomavirus integration in primary oropharyngeal cancers. Genome Res. 32, 55–70 (2022).

10. Pinatti, L. M. et al. Association of human papillomavirus integration with better patient outcomes in oropharyngeal squamous cell carcinoma. Head Neck 43, 544–557 (2021).

11. Lim, M. Y., Dahlstrom, K. R., Sturgis, E. M. & Li, G. Human papillomavirus integration pattern and demographic, clinical, and survival characteristics of patients with oropharyngeal squamous cell carcinoma. Head Neck 38, 1139–1144 (2016).

12. Mainguené, J. et al. Human papilloma virus integration sites and genomic signatures in head and neck squamous cell carcinoma. Mol. Oncol. 16, 3001–3016 (2022).

13. Balaji, H. et al. Causes and Consequences of HPV Integration in Head and Neck Squamous Cell Carcinomas: State of the Art. Cancers 13, (2021).

14. Muñoz-Bello, J. O. et al. Potential Transcript-Based Biomarkers Predicting Clinical Outcomes of HPV-Positive Head and Neck Squamous Cell Carcinoma Patients. Cells 13, (2024).

15. Khanna, S. et al. Determining the molecular landscape and impact on prognosis in HPV-associated head and neck cancer. Cancers of the Head & Neck 5, 1–10 (2020).

16. Münger, K. et al. Mechanisms of Human Papillomavirus-Induced Oncogenesis. J. Virol. 78, 11451 (2004).

17. Graham, S. V. The human papillomavirus replication cycle, and its links to cancer progression: a comprehensive review. Clin. Sci. 131, 2201–2221 (2017).

18. Johansson, C. & Schwartz, S. Regulation of human papillomavirus gene expression by splicing and polyadenylation. Nat. Rev. Microbiol. 11, 239–251 (2013).

19. Schwartz, S., Wu, C. & Kajitani, N. RNA elements that control human papillomavirus mRNA splicing-targets for therapy? J. Med. Virol. 96, e29473 (2024).

20. Graham, S. V. & Faizo, A. A. A. Control of human papillomavirus gene expression by alternative splicing. Virus Res. 231, 83 (2017).

21. Yu, L., Majerciak, V. & Zheng, Z.-M. HPV16 and HPV18 Genome Structure, Expression, and Post-Transcriptional Regulation. Int. J. Mol. Sci. 23, 4943 (2022).

22. Van Doorslaer, K. et al. The Papillomavirus Episteme: a major update to the papillomavirus sequence database. Nucleic Acids Res. 45, D499–D506 (2017).

23. Collins, S. I. et al. Disruption of the E2 gene is a common and early event in the natural history of cervical human papillomavirus infection: a longitudinal cohort study. Cancer Res. 69, 3828–3832 (2009).

24. Sanchez-Perez, A. M., Soriano, S., Clarke, A. R. & Gaston, K. Disruption of the human papillomavirus type 16 E2 gene protects cervical carcinoma cells from E2F-induced apoptosis. J. Gen. Virol. 78 **( Pt** **11**), 3009–3018 (1997).

25. Kamal, M. et al. Human papilloma virus (HPV) integration signature in Cervical Cancer: identification of MACROD2 gene as HPV hot spot integration site. Br. J. Cancer 124, 777–785 (2020).

26. Pinatti, L. M., Walline, H. M. & Carey, T. E. Human Papillomavirus Genome Integration and Head and Neck Cancer. J. Dent. Res. 97, 691–700 (2018).

27. Choo, K. B., Pan, C. C. & Han, S. H. Integration of human papillomavirus type 16 into cellular DNA of cervical carcinoma: preferential deletion of the E2 gene and invariable retention of the long control region and the E6/E7 open reading frames. Virology 161, 259–261 (1987).

28. Cricca, M. et al. Disruption of HPV 16 E1 and E2 genes in precancerous cervical lesions. J. Virol. Methods 158, 180–183 (2009).

29. Xue, Y. et al. Loss of HPV16 E2 Protein Expression Without Disruption of the E2 ORF Correlates with Carcinogenic Progression. Open Virol. J. 6, 163–172 (2012).

30. Gillison, M. L. et al. Human papillomavirus and the landscape of secondary genetic alterations in oral cancers. Genome Res. 29, 1–17 (2019).

31. Singh, A. K. et al. Cis-regulatory effect of HPV integration is constrained by host chromatin architecture in cervical cancers. Mol. Oncol. 18, 1189–1208 (2024).

32. Labarge, B. et al. Human Papillomavirus Integration Strictly Correlates with Global Genome Instability in Head and Neck Cancer. Mol. Cancer Res. 20, 1420–1428 (2022).

33. Warburton, A., Markowitz, T. E., Katz, J. P., Pipas, J. M. & McBride, A. A. Recurrent integration of human papillomavirus genomes at transcriptional regulatory hubs. NPJ Genom Med 6, 101 (2021).

34. Olthof, N. C. et al. Viral load, gene expression and mapping of viral integration sites in HPV16-associated HNSCC cell lines. Int. J. Cancer 136, E207–18 (2015).

35. Zhou, L. et al. Long-read sequencing unveils high-resolution HPV integration and its oncogenic progression in cervical cancer. Nat. Commun. 13, 2563 (2022).

36. Akagi, K. et al. Genome-wide analysis of HPV integration in human cancers reveals recurrent, focal genomic instability. Genome Res. 24, 185–199 (2014).

37. Parfenov, M. et al. Characterization of HPV and host genome interactions in primary head and neck cancers. Proc. Natl. Acad. Sci. U. S. A. 111, 15544–15549 (2014).

38. Fan, J. et al. Multi-omics characterization of silent and productive HPV integration in cervical cancer. Cell Genom 3, 100211 (2023).

39. Choi, Y. J. et al. E2/E6 ratio and L1 immunoreactivity as biomarkers to determine HPV16-positive high-grade squamous intraepithelial lesions (CIN2 and 3) and cervical squamous cell carcinoma. J. Gynecol. Oncol. 29, e38 (2018).

40. Das, P. et al. Human papillomavirus (HPV) genome status & cervical cancer outcome--A retrospective study. Indian J. Med. Res. 142, 525–532 (2015).

41. Ibragimova, M. et al. HPV status and its genomic integration affect survival of patients with cervical cancer. Neoplasma 65, 441–448 (2018).

42. Kiseleva, V. I. et al. The Presence of Human Papillomavirus DNA Integration is Associated with Poor Clinical Results in Patients with Third-Stage Cervical Cancer. Bull. Exp. Biol. Med. 168, 87–91 (2019).

43. Loss of HPV 16-E2 Predicts Disease Free Survival and Promotes Platinum Resistance in Human Papillomavirus-Positive Head and Neck Squamous Cell Carcinoma. International Journal of Radiation Oncology*Biology*Physics 112, e6–e7 (2022).

44. Choi, J.-H. et al. Single-cell transcriptome profiling of the stepwise progression of head and neck cancer. Nat. Commun. 14, 1055 (2023).

45. Puram, S. V. et al. Cellular states are coupled to genomic and viral heterogeneity in HPV-related oropharyngeal carcinoma. Nat. Genet. 55, 640–650 (2023).

46. Bill, R. et al. macrophage polarity identifies a network of cellular programs that control human cancers. Science 381, 515–524 (2023).

47. Rajaby, R. et al. SurVirus: a repeat-aware virus integration caller. Nucleic Acids Res. 49, e33 (2021).

48. Grabherr, M. G. et al. Full-length transcriptome assembly from RNA-Seq data without a reference genome. Nat. Biotechnol. 29, 644–652 (2011).

49. Briggs, A., et al. Nf-Core/Viralintegration: V0.1.1 - Caladrius. (Zenodo, 2023). doi:10.5281/ZENODO.7783480.

50. Huang, S. H. & O’Sullivan, B. Overview of the 8th edition TNM classification for head and neck cancer. Curr. Treat. Options Oncol. 18, 40 (2017).

51. Gel, B. & Serra, E. karyoploteR: an R/Bioconductor package to plot customizable genomes displaying arbitrary data. Bioinformatics 33, 3088–3090 (2017).

52. Szklarczyk, D. et al. STRING v11: protein-protein association networks with increased coverage, supporting functional discovery in genome-wide experimental datasets. Nucleic Acids Res. 47, D607–D613 (2019).

53. Oughtred, R. et al. The BioGRID database: A comprehensive biomedical resource of curated protein, genetic, and chemical interactions. Protein Sci. 30, 187 (2021).

54. Chakravarty, D., et al. OncoKB: A Precision Oncology Knowledge Base. JCO Precis Oncol 2017, (2017).

55. Ggsurvfit. https://www.danieldsjoberg.com/ggsurvfit/.

56. Gray, R. J. Modeling survival data: Extending the cox model. J. Am. Stat. Assoc. 97, 353–354 (2002).

57. Wu, T. et al. clusterProfiler 4.0: A universal enrichment tool for interpreting omics data. Innovation (Camb) 2, 100141 (2021).

58. Leshchiner, I. et al. Inferring early genetic progression in cancers with unobtainable premalignant disease. Nature Cancer 4, 550–563 (2023).

59. Wang, X. et al. Human papillomavirus integration perspective in small cell cervical carcinoma. Nat. Commun. 13, 1–10 (2022).

60. Brant, A. C. et al. Characterization of HPV integration, viral gene expression and E6E7 alternative transcripts by RNA-Seq: A descriptive study in invasive cervical cancer. Genomics 111, 1853–1861 (2019).

61. Santegoets, S. J., Stolk, A., Welters, M. J. P. & van der Burg, S. H. The combined HPV16-E2/E6/E7 T cell response in oropharyngeal cancer predicts superior survival. Cell Reports Medicine 4, (2023).

62. Hibbert, J., Halec, G., Baaken, D., Waterboer, T. & Brenner, N. Sensitivity and Specificity of Human Papillomavirus (HPV) 16 Early Antigen Serology for HPV-Driven Oropharyngeal Cancer: A Systematic Literature Review and Meta-Analysis. Cancers 13, (2021).

63. Kreimer, A. R. et al. Evaluation of human papillomavirus antibodies and risk of subsequent head and neck cancer. J. Clin. Oncol. 31, 2708–2715 (2013).

64. Lang Kuhs, K. A., et al. Human papillomavirus 16 E6 antibodies are sensitive for human papillomavirus-driven oropharyngeal cancer and are associated with recurrence. Cancer 123, 4382–4390 (2017).

65. de Freitas, A. C., de Oliveira, T. H. A., Barros, M. R., Jr & Venuti, A. hrHPV E5 oncoprotein: immune evasion and related immunotherapies. J. Exp. Clin. Cancer Res. 36, 71 (2017).

