## Supplementary methods and figures for "HPV integration in head and neck cancer: downstream splicing events and expression ratios linked with poor outcomes"

**Bulk RNA-seq QC and gene counts calculation**

Quality control of the data was performed using FastQC (v0.11.9), followed by adapter trimming with Cutadapt (v3.4). Reads trimmed were aligned to a custom reference genome, which included hg38 and high-risk HPV sequences (HPV16, HPV18, HPV31, HPV33, HPV35, HPV39, HPV45, HPV51, HPV52, HPV56, HPV58, HPV59, HPV66, HPV68, HPV73, and HPV82), using STAR (v2.7.9a). Human gene expression levels were quantified with htseq-count (v0.13.5) with default settings.

**Calling expressed HPV integration events from bulk and scRNA-seq**

Viral integration detection software designed for paired-end NGS data typically measures the confidence of the viral integrations based on two types of reads: 1) support reads: two reads in one pair with one aligned to the virus genome and the other aligned to the human genome. 2) split reads: a single read that is partially aligned to the human genome with the remaining aligned to the virus genome. Only split reads provide the exact insertion location.

For bulk RNA-seq data, we built the reference genome with hg38 plus high-risk HPV types and employed two integration detection software programs, SurVirus^1^ and CTAT-VIF ^2^. SurVirus uses BWA MEM as the aligner while CTAT-VIF uses STAR. Default settings were applied for both programs across cohorts (UM_FF, UM_FFPE, HVC) except for TCGA. Since TCGA RNA-seq data reads are substantially shorter (48bp vs 100bp minimum for others), we altered the SurVirus argument ‘--minClipSize 5’ to include more soft-clipped reads as evidence in the integration prediction steps.

Since no existing pipeline has been developed to call virus integration based on scRNA-seq data, we modified the nf-core/viralintegration ^3^ pipeline to accomplish this task (VIF-single cell). The same reference files were used to preprocess the bulk data, and we replaced STAR with STAR-solo and modified the intermediate scripts in nf-core/viralintegration to accommodate the format of scRNA-seq bam files. The code for the VIF-single cell pipeline is available at (GitHub link in process). Since the scRNA-seq reads are single-end, the detected events are supported completely by split reads.

**Identification of spliced and low-quality integration events.**

When predicting HPVint events, artifacts generated by specific sequence patterns (such as TATA box) or sequences highly homologous between human and HPV, can potentially result in false-positive predictions. To avoid this, we first removed events that occurred at the same exact base pair (bp) on both the human genome and the HPV genome in more than one patient within the same cohort. These events were highly likely to be false positives caused by sequence patterns or batch effects.

Split reads provide direct evidence of HPV-human genome fusions. To identify confident HPVint events, we required all HPVint events detected by any pipeline (Survirus, CTAT-VIF, or VIF-single cell) to be supported by at least two split reads. Events with fewer than two split reads were considered low-quality events.

Previous studies have indicated that there are highly recurrent integration locations on the HPV genome observed in RNA-seq data likely due to RNA splicing ^4–6^, suggesting that integration events detected via RNA-seq do not precisely predict the actual integration sites at the DNA level. We observed that these highly recurrent HPV locations were at or near known HPV splice junctions using the HPV transcript maps, ^7^ and defined these events as ‘spliced’ integration events for further analysis (HPV16 (226-230, 406-408, 880-883, 1303, 2705-2708, 3354-3357, 4179-4181, 5635-5638, 7166), HPV35 (883, 884), HPV33 (894, 233) HPV18 (930)).

**Additional steps to define sample/participant HPV integration status**

Due to RNA degradation and fragmentation in FFPE samples ^8^, SurVirus and CTAT-VIF filtered out some true positive integration events from UM_FFPE samples, due to either the requirement for sequencing depth in integration sites or the insert size of paired reads. Therefore, when judging the UM_FFPE samples' integration status, we manually retrieved HPV integration events filtered out for these reasons by either CTAT-VIF or SurVirus (Supplementary Figure S1B).

HPV integration can cause the loss of E2 ^9^ or other genes (E1, E4, E5, L1 or L2) whose expression is required and tightly regulated in the episomal (non-integrated) state. Thus, we reassigned patients to the HPVint(+) group if they lost expression of any of those HPV genes. To account for varying sequence depth and total HPV expression, for each non-oncogenic gene *G_i_* we calculated:

$$Log2(\frac{Gi}{(E6+E7+1)})$$

We then calculated the z-score of this ratio (HPV gene ratio score) for each cohort of HPVint(-) and HPV-ND samples to define the expected distribution of expression. Samples with z-score smaller than -3 were reassigned to HPVint(+) group.

**Annotation of HPV integration events**

To study the impact of the detected HPV integration events, we used three methods to assign the insertion sites to potentially affected human genes: 1) the *annotatr* Bioconductor package ^10^, 2) assigning the site to the gene with the nearest transcription start site (TSS) or by known enhancer-gene links with the *chipenrich* Bioconductor package (chipenrich.data locusdef.hg38.enhancer_plus5kb) ^11^, and 3) determining if the site is inside a UCSC genome browser-defined gene region using AnnotationHub ^12^. All of the resulting annotations, including gene symbols and genomic region information, were retained for downstream analysis.

Because an integration event can impact human genes other than the nearest gene or known enhancer-gene link, we also identified the two nearest flanking genes on either side of the integration assigned gene (2 upstream and 2 downstream, named left2, left1, right1, and right2), using chipenrich.data locusdef.hg38.nearest.gene. We defined those genes as ‘nearby_genes’.

As various methods can yield different annotations for each event, we manually selected the most appropriate annotation based on a hierarchy reflecting the relevance of each region type to gene expression: 'genes_firstexons '> 'genes_exons' > 'genes_introns' > 'genes_5UTRs' > 'genes_promoters' > 'genes_1to5kb' > 'genes_3UTRs' > 'genes_intronexonboundaries' > 'genes_exonintronboundaries' > 'genes_cds' (from annotatr) > 'enhancer_or_nearest_TSS' (from chipenrich.data) > 'inside_genes' > (From AnnotationHub) > ‘nearby_genes’.

**Identifying HPV integration events linked to human genes with outlier expression**

We normalized the raw gene count data from all bulk RNA-seq data by sequencing depth using the log counts per million metric (logCPM) with the cpm function from the edgeR (3.34.1) Bioconductor package. We calculated z-scores for each gene across samples from each cohort (UM_FF, UM_FFPE, HVC, TCGA). A gene-sample pair was considered an HPV integration-induced expression outlier if its expression in the integrated sample had a z-score absolute value exceeding 2.0.

Since each HPV integration event could be assigned to multiple genes, for heatmap visualization, we selected the assigned gene from each event with the largest z-score absolute value as the “integrated genes”. Although this biases the ‘integrated genes’ to have higher absolute z-scores, it facilitates identification of the most likely target gene of the integration event.

**Calculation of HPV gene features and correlations**

Reads mapped to the relevant high-risk HPV genome were extracted from bam files using Samtools for both bulk and scRNA-seq. For each HPV gene, we summarized all reads starting within it as the raw expression count. We only considered reads whose mates were reverse-strand. We normalized the HPV gene count to log2CPM values with human library size and a prior.count 0.25 the keep consistent with cpm function in edgeR for downstream analysis. We used Pearson correlation coefficients to calculate the correlations among the log2CPM values of HPV genes E1, E2, E5, E6, E7, L1, and L2 and visualized the correlation matrix using the ComplexHeatmap package. Correlations were calculated and visualized separately for each RNA-seq cohort (UM_FF, UM_FFPE, TCGA, HVC, and single-cell) as well as for all samples together. We selected the HPV gene pairs with the highest correlation (L1-L2, E2-E5, E6-E7) to define the short ImmunOnco score (see methods below).

**Calculation and transformation of cell type proportions**

To perform cell type deconvolution across the 236 bulk RNA-seq samples from four cohorts (UM_FF, UM_FFPE, HVC, and TCGA), we first adjusted for batch effects in the human gene counts using Combat-seq ^13^. We then calculated the TPM values for these samples and performed cell type deconvolution using CIBERSORTx with the provided single-cell RNA-seq HNSCC matrix file ^14^. The Impute Cell Fractions Analysis Module and B mode batch correction were selected, and quantile normalization and absolute mode were disabled as recommended. The number of permutations was set to 100. Cell type deconvolution results yielded 10 cell type proportions: CD4+ T cells, CD8+ T cells, B cells, malignant cells, macrophages, fibroblasts, myocytes, endothelial cells, mast cells, and dendritic cells.

Cell types with over 15% zeros were converted to binary variables based on values above or below the median, because their distributions could not be approximated with a normal distribution. CD8+ T cells, B cells, mast cells, and myocytes were transformed using this method, and logistic regression was performed for these cell types using the cell type proportion category as the dependent variable. For cell types with less than 15% zeros, cell type proportions exhibiting high skewness were log-transformed to approximate a normal distribution, which included CD4+ T cells, fibroblasts, dendritic cells, and endothelial cells. The malignant cells, macrophages, and total T cells were not transformed as their proportion distributions were approximately normally distributed.

**Definition of ‘combined’ and ‘random’ sites for DNA vs RNA HPV integration sites comparison.**

The ‘combined’ sites were defined as the union set of confident and spliced HPV integration sites. To generate random genomic sites, we used hg38 chromosome sizes to define appropriate ranges for random positioning. A total of 800 random sites were created by sampling chromosomes and assigning a random position within each selected chromosome’s length. These ‘random’ sites were then compared to a reference set of WGS sites to calculate the nearest distance between each random site and its closest WGS site, which allowed us to assess proximity relationships for randomly distributed genomic positions relative to actual WGS sites.

**Definition of productive and silent HPV integration events.**

The positions of virus-host DNA breakpoints from the previous study (David saymer) ^4^ were based on hg19. To align these with our RNA-seq data, we employed the liftOver tool (version 1.28.0) in R to convert from hg19 to hg38. To merge nearby integration sites in a sample, those within 10kb of each other and annotated to the same gene were consolidated into a single integration event due to their proximity and identical target gene. A DNA integration event was classified as ‘productive’ if an RNA event was found within ±100kb in the same participant. The remaining DNA events were defined as ‘silent’. To annotate genes at each DNA and RNA breakpoint, we utilized the hierarchical system described above.

**Supplementary Figure Legends.**

**Supplementary Figure S1. Workflow for determining the HPV integration status for RNA-seq samples.** (A) General workflow applicable to both FF and FFPE samples (B) Specific workflow for UM_FFPE samples. (C) Additional steps using the HPV gene ratio to correct samples misclassified as HPVint(-) or HPVint-ND. The solid line represents the sample is reclassified to HPVint(+).

**Supplementary Figure S2. Histograms of cell deconvolution results from 8 cell types.**

**Supplementary Figure S3.** (A) Venn diagram comparing our identified HNSCC recurrent sites with CC recurrent sites from a previous paper. (B) Z-score heatmap for all sample-gene pairs related to non-recurrent genes.

**Supplementary Figure S4. (**A) Correlation heatmaps of HPV gene log2cpm for each cohort. (B-C) Violin plots across cohorts illustrating the ImmunOnco score across (B) E1* integration, Not E1* integration, and HPVint(-) groups (C) E6* integration, Not E6* integration and HPVint(-) groups.

**Supplementary Figure S5. Extra survival tests for HPV integration status, E1* integration status and ImmunOnco score.** (A-B) Overall survival KM and Forest plot for HPVint(+) versus HPVint(-),Forest plot with AJCC v8 stage, age, and smoking status as covariates.(C) Forest plot compare the E1* integration versus HPVint(-). (D-G) Overall survival KM and Forest plots for UM_FFPE and FF samples (UM_FF+TCGA) ImmunOnco score, Forest plot with AJCC v8 stage, age, and smoking status as covariates and KM plot is based on tertiles.

**Supplementary Figure S6. Extra recurrence tests for HPV integration status, E1* integration status, ImmunOnco, short ImmunOnco score and additional features.** (A-B) recurrence KM and Forest plots for Short ImmunOnco score using UM_FFPE samples, Forest plot with AJCC v8 stage and age as covariates. (C-E) Recurrence KM plots for E1/(E6+E7),(E2+E5)/(E6+E7) and (L1+L2)/(E6+E7) using UM_FFPE samples. (F-G) Recurrence Forest plots for log2(E2/E6), E6 log2CPM and E7 log2CPM using UM_FFPE samples, Forest plots with AJCC v8 stage and age as covariates.

**Supplementary Figure S7.** (**A**) Heatmap of Z-score illustrating DEGs, showing the 220 top DEGs (FDR < 0.05) between double-positive patients (orange) and double-negative patients (dark blue). HPV integration status, cohorts, sex, molecular subtypes, and adjusted ImmunOnco scores are annotated at the top. (**B**) PCA plot for the three categories of integration, using the 220 top DEGs.

**Supplementary Table Descriptions.**

**Supplementary Table S1**

S1.1 All HPV integration sites detected with their assigned genes and genomic annotations.

S1.2 Unique unspliced sites sample-chr-position pairs

S1.3 Unique spliced sites sample-chr-position pairs

S1.4 All genes involved in those HPV integration sites.

**Supplementary Table S2.** **HPV integration status for all samples.**

**Supplementary Table S3. All patients collected demographic information**

**Supplementary Table S4**

S4.1 Recurrent gene clusters defined

S4.2 HNSCC unique recurrent gene clusters not captured by CC.

S4.3 Integrated expression outlier genes have more than 3 protein interactions.

**Supplementary Table S5**

S5.1 Samples level ImmunOnco and short ImmunOnco scores

S5.2 Patients level ImmunOnco and short ImmunOnco scores

S5.2 Patients Survival status

S5.3 Patients Recurrent status

**Supplementary Table S6. DEGs with consistent directions across cohorts by different comparisons**

S6.1 HPVint(+) versus HPVint(-)

S6.2 high versus low Immunonco score

S6.3 E1* integration versus not E1* integration for HPVint(+) patients only

**Supplementary Table S7. Gene sets representative analysis across cohorts for different comparisons**

S7.1 HPVint(+) to HPVint(-)

S7.2 high to low Immunonco score

S7.3 E1* integration or not E1* integration

**Supplementary Table S8. linear regression or ANOVA analysis for cell deconvolution results versus HPV integration status, E1* integration or ImmunOnco score.**

S8.1 HPVint(+) to HPVint(-)

S8.2 E1* integration

S8.2 Immunonco score

**Supplementary Table S9. 102 patients DNA and RNA level HPV integration status**

**Supplementary Table S10. 102 patients identified HPV integration sites**

S8.1 DNA level

S8.2 RNA level

**Supplementary Table S11. Significant DEGs identified between double positive and double negative patients.**

**Supplemental References**

1.	Rajaby, R. *et al.* SurVirus: a repeat-aware virus integration caller. *Nucleic Acids Res.* **49**, e33 (2021).

2.	Grabherr, M. G. *et al.* Full-length transcriptome assembly from RNA-Seq data without a reference genome. *Nat. Biotechnol.* **29**, 644–652 (2011).

3.	Briggs, A. *et al.* *Nf-Core/Viralintegration: V0.1.1 - Caladrius*. (Zenodo, 2023). doi:10.5281/ZENODO.8164980.

4.	Symer, D. E. *et al.* Diverse tumorigenic consequences of human papillomavirus integration in primary oropharyngeal cancers. *Genome Res.* **32**, 55–70 (2022).

5.	Fan, J. *et al.* Multi-omics characterization of silent and productive HPV integration in cervical cancer. *Cell Genom* **3**, 100211 (2023).

6.	Brant, A. C. *et al.* Characterization of HPV integration, viral gene expression and E6E7 alternative transcripts by RNA-Seq: A descriptive study in invasive cervical cancer. *Genomics* **111**, 1853–1861 (2019).

7.	Yu, L., Majerciak, V. & Zheng, Z.-M. Correction: Yu et al. HPV16 and HPV18 Genome Structure, Expression, and Post-Transcriptional Regulation. 2022, , 4943. *Int. J. Mol. Sci.* **23**, (2022).

8.	Newton, Y. *et al.* Large scale, robust, and accurate whole transcriptome profiling from clinical formalin-fixed paraffin-embedded samples. *Sci. Rep.* **10**, 17597 (2020).

9.	Nishimura, A. *et al.* Mechanisms of human papillomavirus E2-mediated repression of viral oncogene expression and cervical cancer cell growth inhibition. *J. Virol.* **74**, 3752–3760 (2000).

10.	Cavalcante, R. G. & Sartor, M. A. annotatr: genomic regions in context. *Bioinformatics* **33**, 2381–2383 (2017).

11.	Welch, R. P. *et al.* ChIP-Enrich: gene set enrichment testing for ChIP-seq data. *Nucleic Acids Res.* **42**, e105–e105 (2014).

12.	Morgan, M. & Shepherd, L. *AnnotationHub: Client to Access AnnotationHub Resources*. (2024).

13.	Zhang, Y., Parmigiani, G. & Johnson, W. E. : batch effect adjustment for RNA-seq count data. *NAR Genom Bioinform* **2**, lqaa078 (2020).

14.	Newman, A. M. *et al.* Determining cell type abundance and expression from bulk tissues with digital cytometry. *Nat. Biotechnol.* **37**, 773–782 (2019).

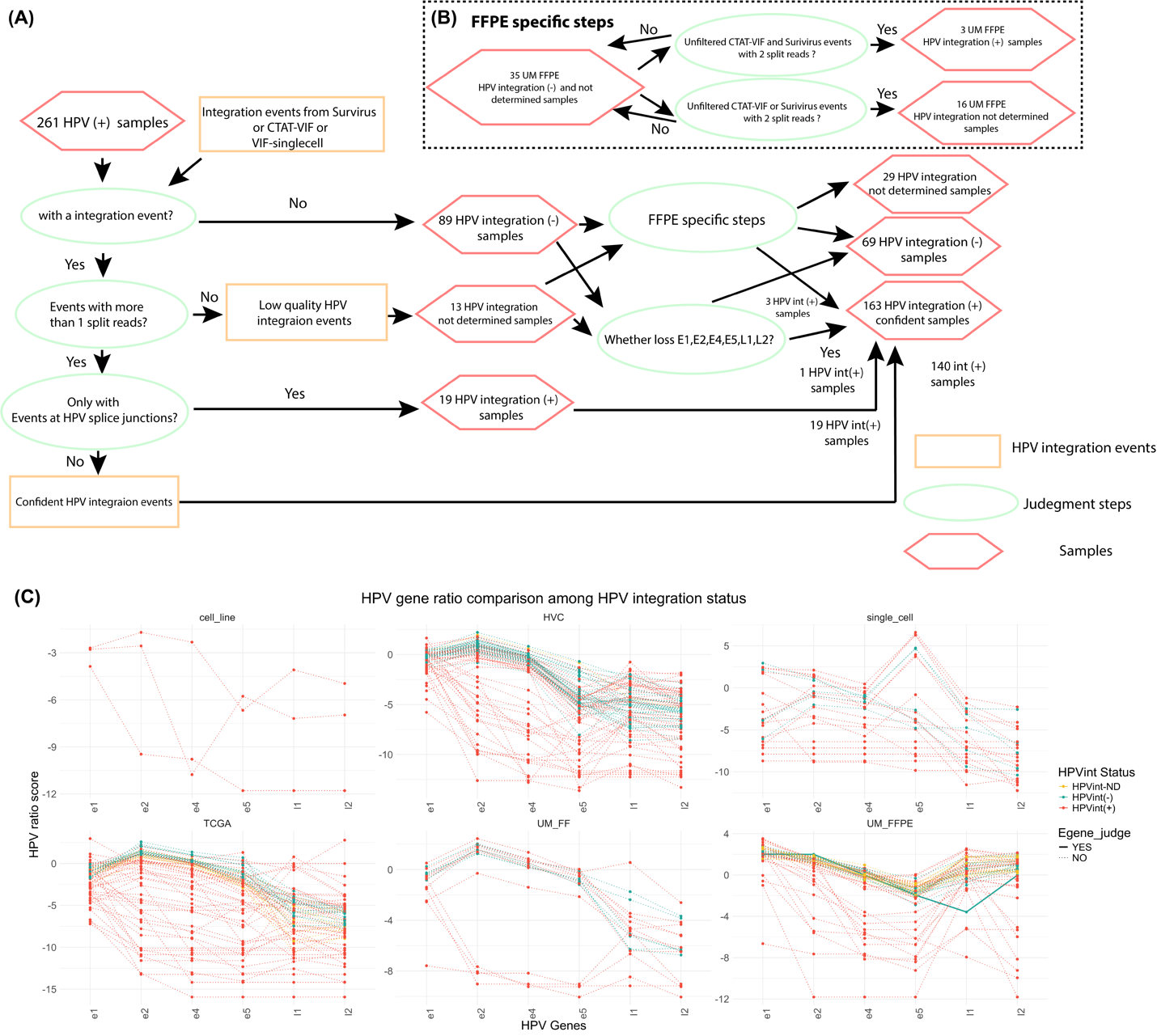

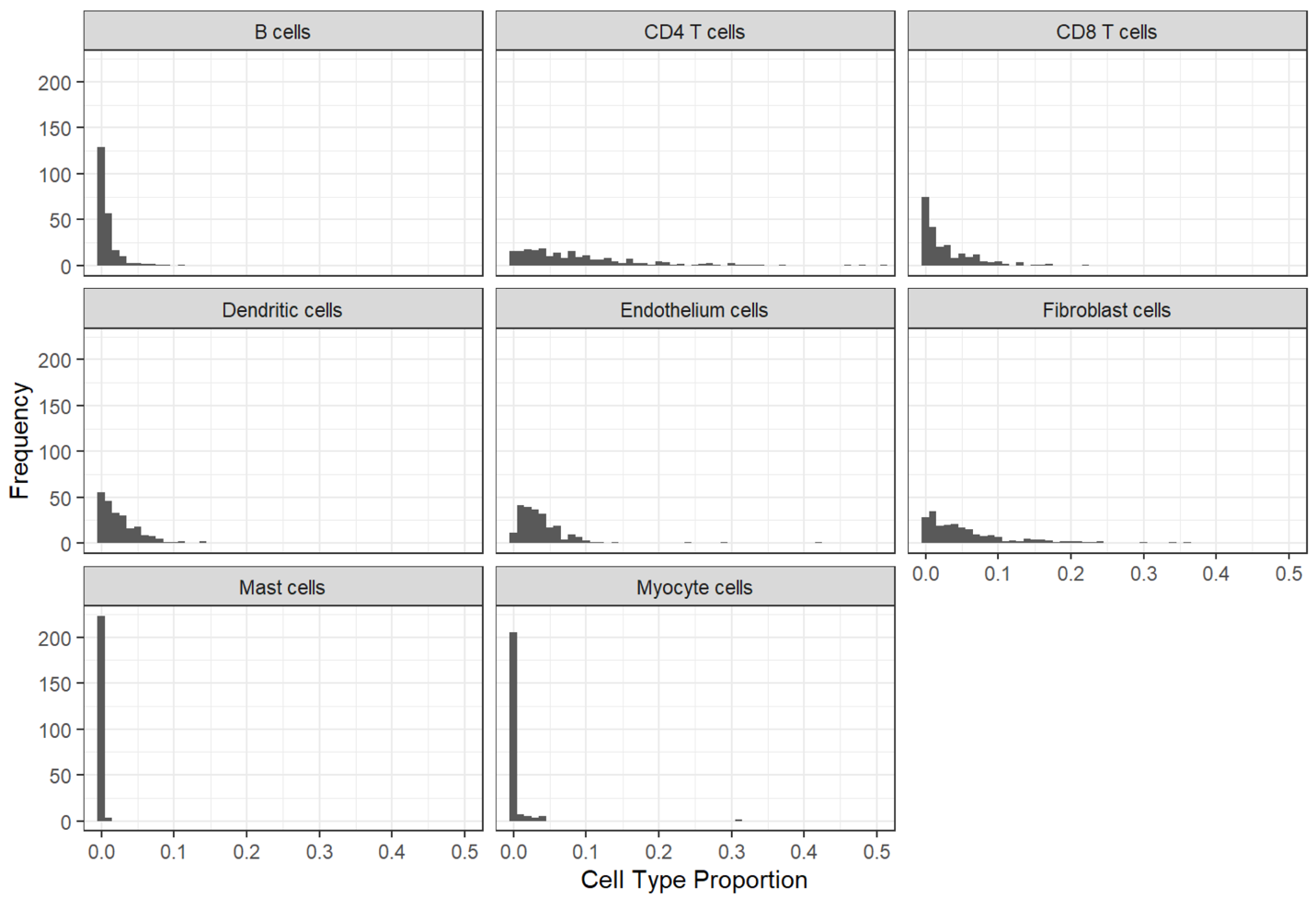

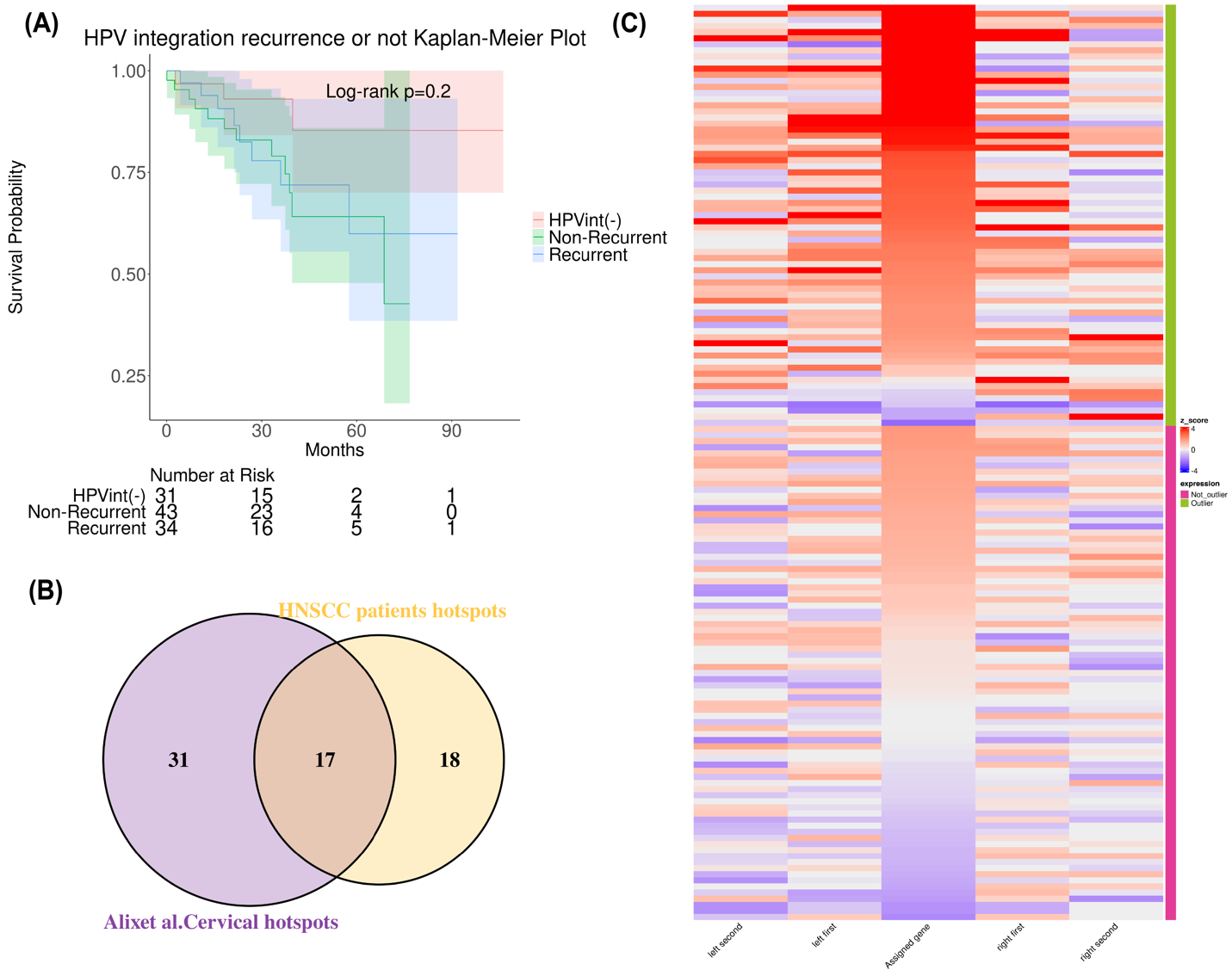

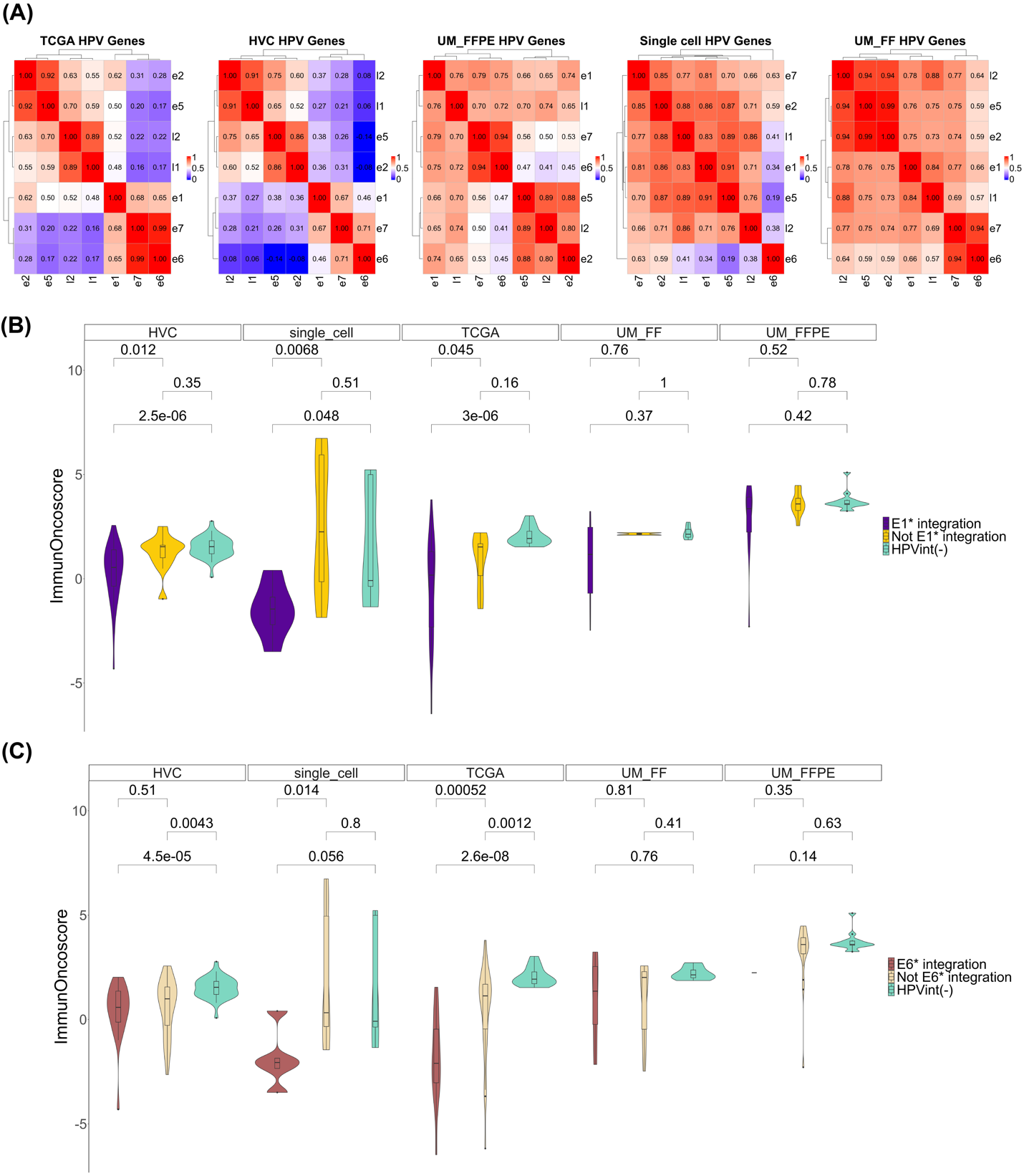

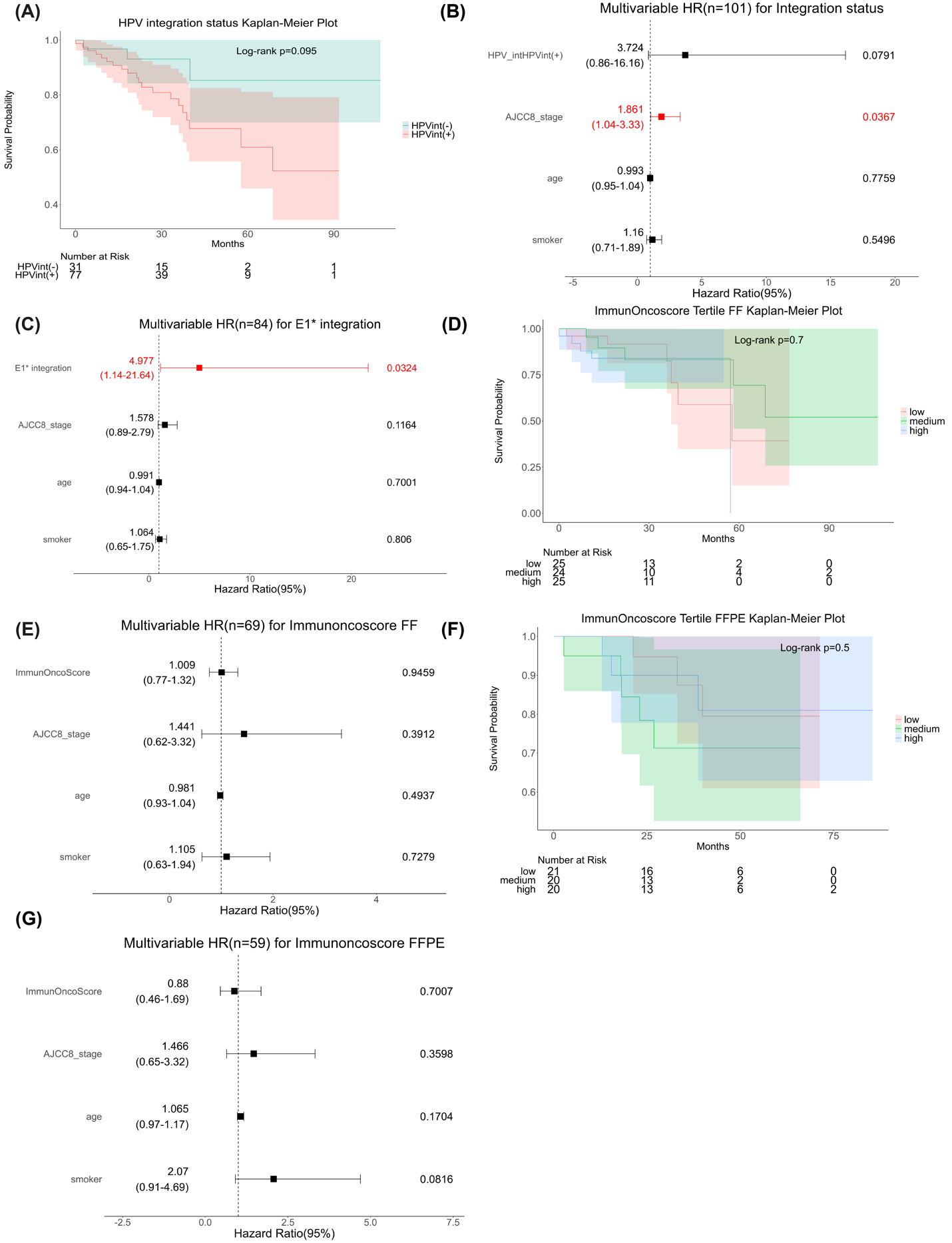

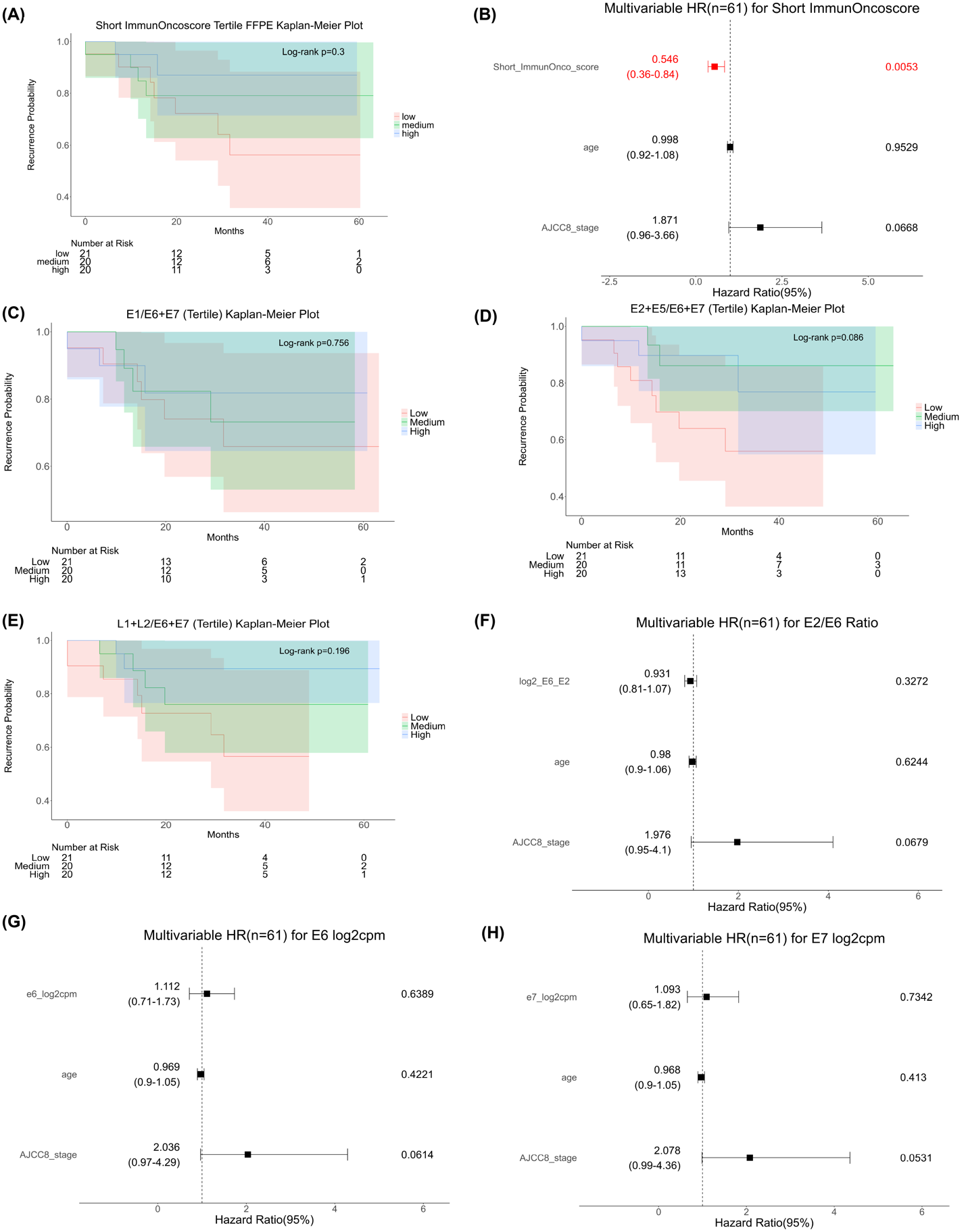

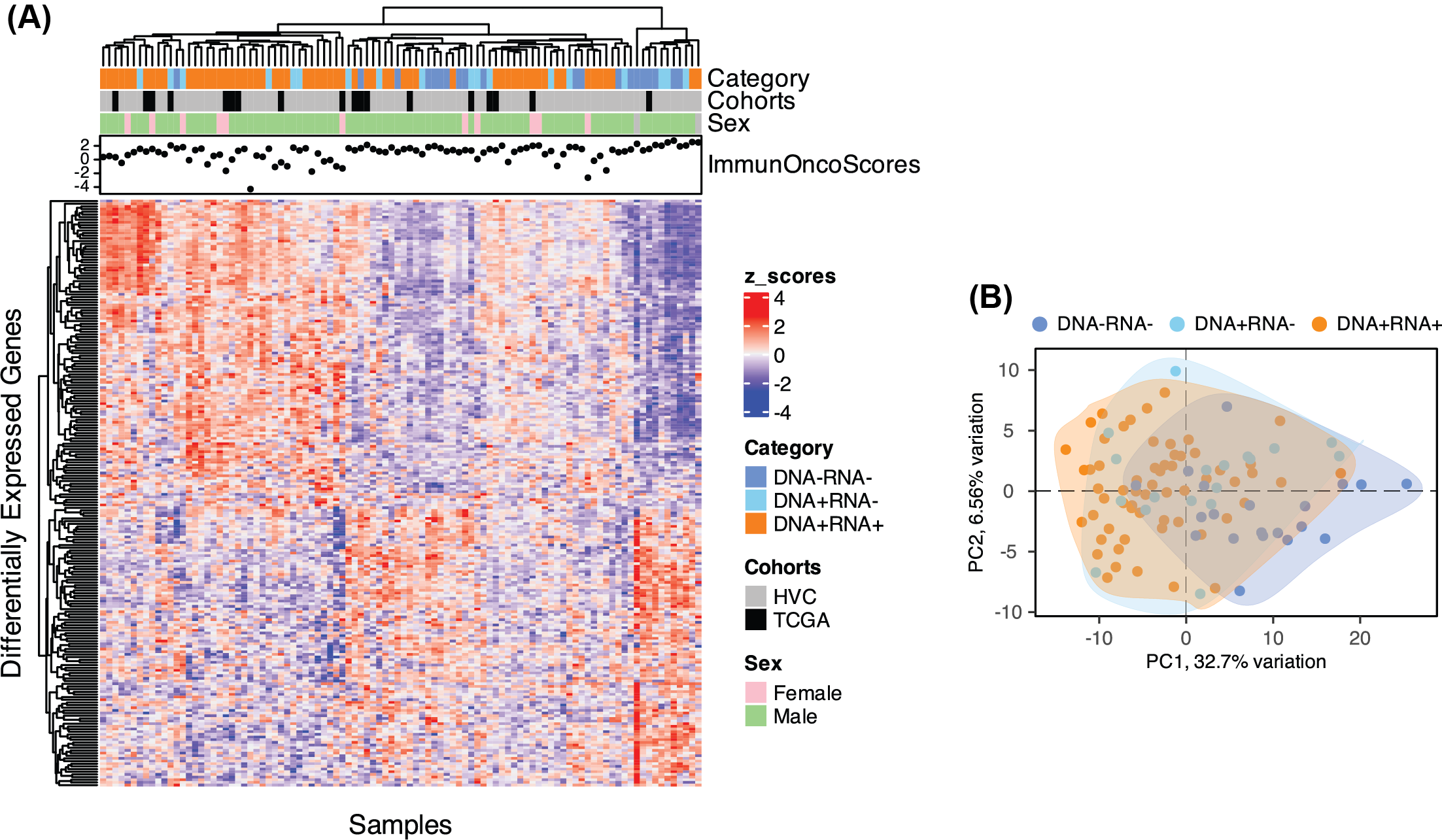
